# Xenon improves autism-like behaviors in mice through potentiation of PVT–CeA glutamatergic neurotransmission

**DOI:** 10.64898/2026.09.23.753714

**Authors:** Jinpiao Zhu, Yina Ruan, Juntao Weng, Xiao Xu, Ziang Wu, Yonglin Yu, Haifeng Li, Xuekun Li, Jianbin Tong, Wen Ouyang, Qiang Shu, Daqing Ma

**Author notes:** These authors contributed equally to this work (Jinpiao Zhu, Yina Ruan, Juntao Weng).

## Abstract

Autism spectrum disorder (ASD) is a common neurodevelopmental disease with no currently effective treatments. In this study, Xenon (Xe), a noble gas used clinically as an anesthetic and a neuroprotectant, has been investigated for its potential therapeutic effects in ASD. Two genetic ASD mouse models (*Mef2c* mutant and *Shank3^−/−^*) were exposed to a low-concentration of xenon (25%) mixed with oxygen (30%) and balanced with nitrogen for a single exposure of 30 min or for 10 min per day for 10 days. These Xe exposure paradigms rescued ASD-like behaviors in both male and female mice. The behavioral improvements persisted for up to 10 days following termination of the 10-day exposure paradigm. Mechanistically, Xe reduced glutamic acid decarboxylase expression in the inhibitory neurons (verified with RNAseq) of the paraventricular thalamic nucleus (PVT) and decreased GABA release during social interaction. Targeted infusion the GABA_A_ receptor agonist muscimol into the PVT abolished the Xe-mediated behavioral improvements. Reduced GABAergic transmission following Xe exposure increased PVT glutamatergic neuronal activity. Chemogenetic inhibition of these neurons eliminated the therapeutic effects of Xe, whereas chemogenetic activation mimicked the Xe-induced behavioral improvements. Furthermore, opto-inhibition of PVT glutamatergic projections to the central amygdala (CeA) blocked Xe-mediated behavioral improvement. Opto-activation of the PVT–CeA glutamatergic circuit reversed ASD-like behaviors. These findings demonstrated that Xe ameliorates ASD-like phenotypes in mouse models by modulating the PVT^Glu^–CeA circuit, supporting Xe as a novel potential therapy for ASD.

## INTRODUCTION

Autism spectrum disorder (ASD) is a common neurodevelopmental illness characterized by core features of social communication impairment, and restricted and repetitive behaviors. Recent epidemiological data from the United States indicate that ASD affects approximately 1 in 31 individuals(*1*), and the global prevalence is estimated to exceed 61.8 million cases(*2*).

ASD is now recognized as one of the top ten causes of non-fatal health burden among individuals under 20 years of age(*2*). The mechanisms underlying ASD remain incompletely understood, but converging evidence indicates that neural circuit dysfunction plays a central role in its pathogenesis. Targeting specific brain regions and their associated circuits has been shown to ameliorate ASD-like symptoms in human(*3, 4*) and animal models(*5, 6*). The paraventricular thalamic nucleus (PVT), a key thalamic hub, is involved in social behavioral and emotional regulation(*7*). The PVT sends dense projections to the nucleus accumbens (NAc), and selective inhibition of the PVT–NAc circuit exacerbates stress-induced social deficits, whereas activation of this circuit ameliorates these deficits(*8*) and promotes prosocial rescue-like behaviors(*9*). Recent immunohistochemical analyses reported a reduction in nicotinic acetylcholine receptor–positive neurons in the PVT of individuals with ASD, suggesting that region-specific neurobiological alterations may be involved in ASD development(*10*). Collectively, these findings highlight the PVT as a critical node in ASD-related neural circuitry and suggest that targeting PVT pathways may offer a promising therapeutic strategy for ASD.

Existing pharmacologic treatments, *e.g.,* aripiprazole and risperidone, primarily target co-occurring behavioral and emotional dysregulation but their effects are inadequate(*11, 12*). Therefore, there remains a critical need for therapeutic strategies capable of minimizing or attenuating ASD symptoms. Multiple medical gases are safely used in clinical practice, including general anesthesia and hyperbaric oxygen therapy. Xenon (Xe), a noble gas with a fully filled valence shell, is chemically inert and has been used as a general anesthetic since the mid-20th century(*13*). Of note, a growing body of *in vivo* and *in vitro* studies demonstrated robust neuroprotective effects of Xe in models of traumatic brain injury and neuronal damage(*14–17*). Although the precise molecular mechanisms of Xe action remain incompletely defined, its actions on glutamatergic neurotransmission are among the most well characterized(*18, 19*). More recently, inhaled Xe has been shown to mitigate pathological changes in mouse models of amyloidosis and tauopathy by shifting microglial states toward a pre-neurodegenerative, homeostatic phenotype *via* interferon-γ signaling(*20*). In agreement with these findings, our pilot study using a valproic acid (VPA)-induced rat ASD model demonstrated that acute Xe inhalation improved social behavior, decreased abnormal exploratory activity, and restored normal performance in VPA-induced autism rats; however, the underlying mechanisms remain unclear (*21*). Given that genetic mutation represents a major contributor to ASD pathogenesis, it is unknown whether Xe inhalation can alleviate autism-like phenotypes in genetic ASD models and whether such effects are mediated through modulation of the PVT glutamatergic system.

In the present study, we investigated the therapeutic potential of Xe inhalation in ASD using multiple genetic mouse models, including *Mef2c* mutant (c.104T>C, p.L35P; L35P^+/−^) mice (*22*) and *Shank3* knockout (*Shank3^−/−^*) mice. The L35P^+/−^ mice was used to represent deficits in neuronal formation and differentiation, whereas *Shank3^−/−^* mice captured impairments in synaptic development and plasticity. We found that Xe inhalation reduced the expression of glutamate decarboxylase, leading to decreased GABA synthesis and release. This consequently enhanced PVT glutamatergic neuronal activity. Of note, Xe selectively increased the activity of PVT glutamatergic neurons projecting to the central amygdala (CeA), thereby rescuing deficits in ASD-like behaviors. Collectively, these findings suggest that Xe restores molecular, cellular and circuit-level dysregulation in ASD and may be a promising therapy for ameliorating ASD phenotypes.

## RESULTS

### Xe improved autism-like behaviors in mice

To investigate whether xenon exposure improves autism-like behaviors, two genetic mouse models of ASD including *Mef2c* mutant mice (L35P^+/−^) and *Shank3* knockout mice (*Shank3^−/−^*) were used (Fig. 1A, and fig. S1A, S2A, and S3A). Mice were exposed to 25% Xe for 10 min per day over 10 consecutive days. Behavioral assessments including the novel object recognition test (NORT), three-chamber social test (TCST) and Y-maze test (YMT), were conducted to evaluate behavioral changes at baseline (day 0), on day 13 (2 days after termination of Xe exposure), and on day 20 (10 days after termination of Xe exposure) (Fig. 1B, and fig. S1B, S2B, and S3B). Compared with wild-type (WT) male mice, L35P^+/−^ male mice exhibited a decreased discrimination index in the NORT (0.43 ± 0.04 vs. 0.58 ± 0.02 in controls, p < 0.01; Fig. 1C), a decreased social preference index in the TCST (0.47 ± 0.03 vs. 0.60 ± 0.02 in controls, p < 0.01; Fig. 1D), and a decreased novel arm preference index in the YMT (0.53 ± 0.03 vs. 0.69 ± 0.04 in controls, p < 0.01; Fig. 1E). Of note, 10-day Xe exposure rescued these behavioral deficits, as evidenced by increased discrimination index (0.58 ± 0.02 vs. 0.49 ± 0.03 in L35P^+/−^ controls, p < 0.05; Fig. 1F), social preference index (0.60 ± 0.03 vs. 0.48 ± 0.04 in L35P^+/−^ controls, p < 0.05; Fig. 1G), and novel arm preference index (0.69 ± 0.03 vs. 0.47 ± 0.03 in L35P^+/−^ controls, p < 0.01; Fig. 1H) on day 13. Furthermore, after a 10-day recovery period without Xe exposure, the therapeutic effects were sustained and remained evident on day 20 (Fig. 1I–K). Xe also rescued behavioral deficits in female L35P^+/−^ mice on both day 13 and day 20, as demonstrated by increased discrimination index, social preference index, and novel arm preference index (fig. S1A-J). Furthermore, we examined whether short-term xenon exposure can attenuate autism-like behaviors (fig. S2A). A single 30-min Xe exposure was sufficient to increase discrimination index, social preference index, and novel arm preference index in both male (fig. S2B-D) and female L35P^+/−^ mice (fig. S2E-G).

**Fig. 1.**
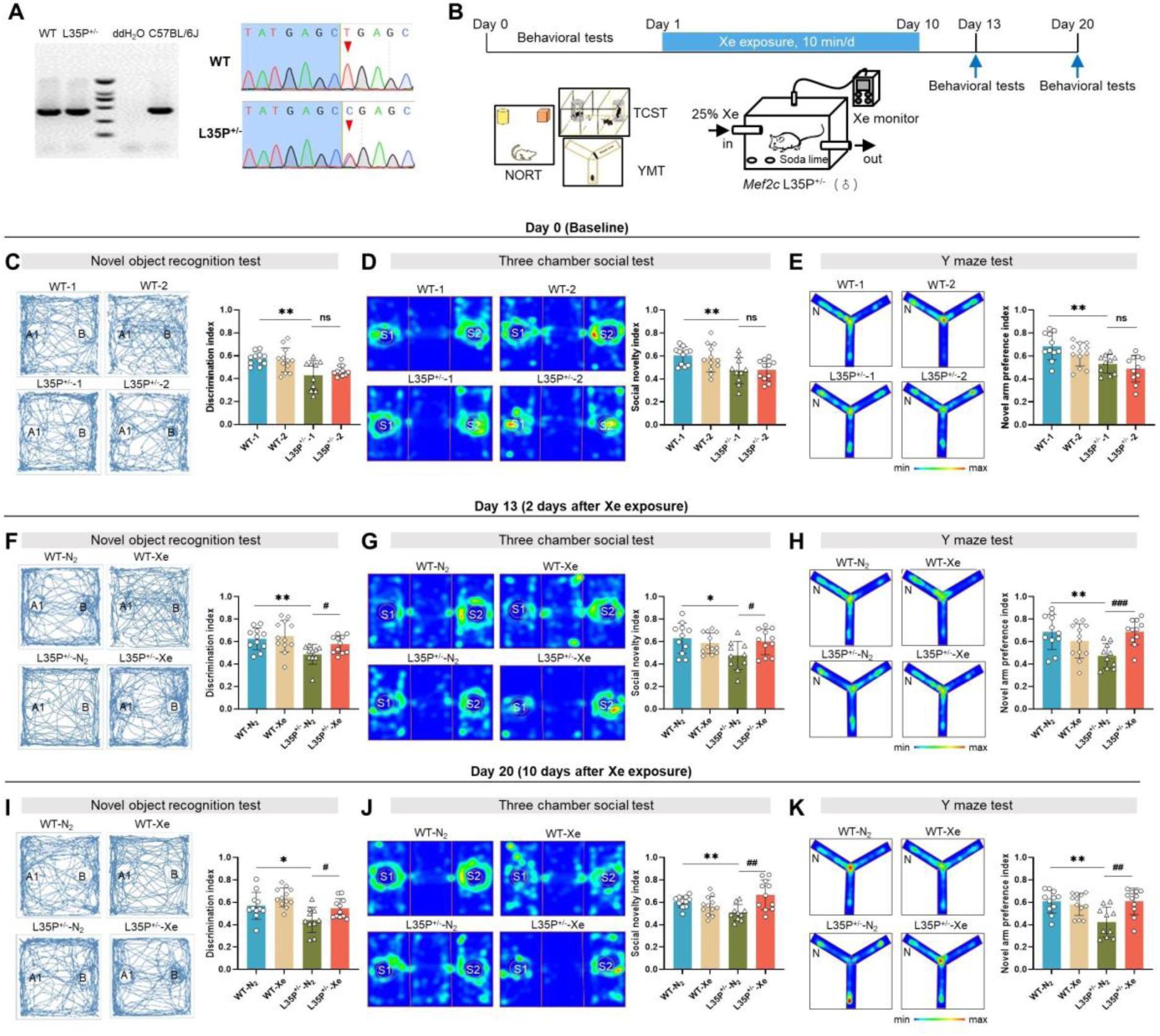
Xe improved autism-like behaviors in male L35P^+/-^ mice. **(A)** Genotyping (left) and Sanger sequencing (right) confirming wild-type (WT) and L35P^+/-^ mice. **(B)** Experiment timeline of xenon (Xe) exposure and behavioral tests including novel object recognition test (NORT), three-chamber social test (TCST) and Y-maze test (YMT). **(C–E)** Baseline behavioral performance in WT and L35P^+/-^ mice without Xe exposure. Representative movement traces or heatmaps (left) and quantification of discrimination index in NORT **(C)**, social novelty index in TCST **(D)**, and novel arm preference index in YMT **(E)**. **(F–H)** Behavioral performance 2 days after Xe exposure. Representative movement traces or heatmaps (left) and quantification of discrimination index in NORT **(F)**, social novelty index in TCST **(G)**, and novel arm preference index in YMT **(H)**. **(I–K)** Behavioral performance 10 days after Xe exposure. Representative traces or heatmaps (left) and quantification of discrimination index in NORT **(I)**, social novelty index in TCST **(J)**, and novel arm preference index in YMT **(K)**. Data are presented as mean ± SEM. \**P* < 0.05, \*\**P* < 0.01, and \*\*\**P* < 0.001; ^#^*P* < 0.05, ^##^*P* < 0.01.

To determine whether the therapeutic effects of Xe are generalizable across distinct genetic ASD models, we next evaluated its effects in *Shank3^−/−^* mice. Compared with WT mice, male *Shank3^−/−^* mice exhibited impairments in cognitive and social behaviors, as shown by a reduced discrimination index in the NORT (0.44 ± 0.02 vs. 0.56 ± 0.04 in controls, p < 0.05; fig. S3C), a decreased social preference index in the TCST (0.51 ± 0.04 vs. 0.64 ± 0.05 in controls, p < 0.05; fig. S3D) and a reduced novel arm preference index in the YMT (0.53 ± 0.04 vs. 0.75 ± 0.04 in controls, p < 0.001; fig. S3E). Of note, 10-day xenon exposure rescued these behavioral deficits in male *Shank3^−/−^* mice, as demonstrated by increases in discrimination index (0.59 ± 0.05 vs. 0.56 ± 0.05 in controls, p > 0.05; fig. S3F), social preference index (0.58 ± 0.03 vs. 0.65 ± 0.04 in controls, p > 0.05; fig. S3G), and novel arm preference index (0.56 ± 0.07 vs. 0.68 ± 0.04 in controls, p > 0.05; fig. S3H) on day 13 compared with controls. Moreover, these improvements were sustained after a 10-day recovery period, with therapeutic effects remaining evident on day 10 after exposure (fig. S3I–K). Consistent with findings in the L35P^+/−^ model, Xe exposure also ameliorated autism-like behavioral deficits in female *Shank3^−/−^* mice on both day 13 and day 20, as shown by improved performance in the NORT, TCST, and YMT compared with controls (Fig. S4B–K).

### PVT glutamatergic neurons were critical for Xe-mediated improvements in autism-like behaviors

Next we investigated which brain regions can be activated by Xe-exposure. C57/BL6J mice were subjected to a 30-min Xe exposure followed by a 30-min recovery period and then processed for whole-brain c-Fos staining (fig. S5A). Compared to controls, Xe significantly increased c-Fos expression in multiple brain regions including the frontal association cortex (FrA), prelimbic cortex (PrL), intermediate part of the lateral leptal nucleus (LSI), paraventricular thalamic nucleus (PVT), paraventricular hypothalamic nucleus (PVN), and central amygdaloid nucleus (CeA) (fig. S5B-C). To further determine whether Xe activates specific neuronal populations in our mouse autism model, L35P^+/−^ mice were also exposed to Xe followed by c-Fos staining. Of note, Xe exposure increased c-Fos expression in the PVT, including the anterior (PVA), medial (PVM), and posterior (PVP) subdivisions (fig. S5D-E).

Moreover, the majority of Xe-induced c-Fos–positive cells in the PVT were co-labeled with calcium/calmodulin-dependent protein kinase II subunit alpha (CaMKII α), a marker for excitatory glutamatergic neurons, indicating selective activation of PVT glutamatergic neurons by Xe (Fig. 2A-B).

**Fig. 2.**
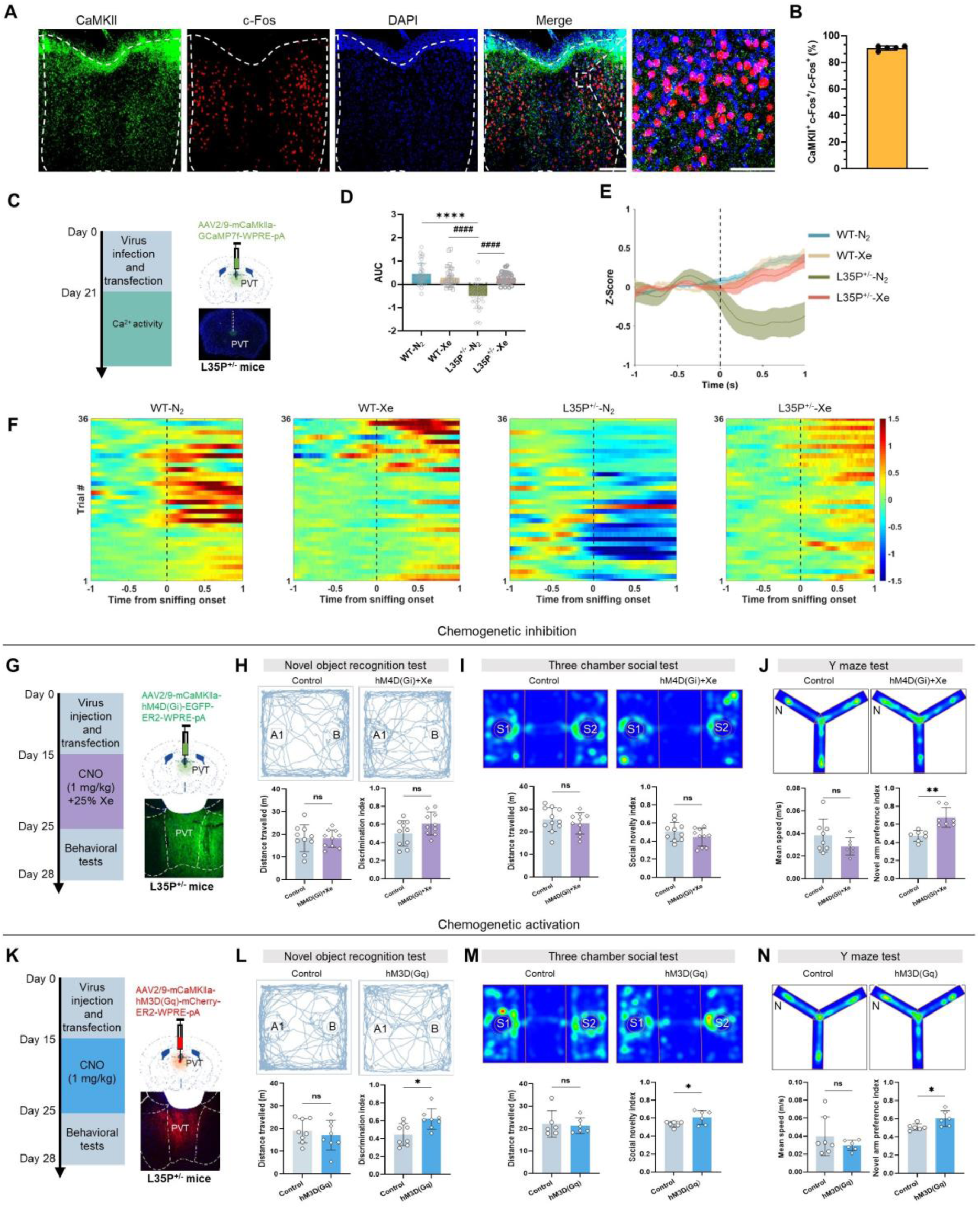
PVT glutamatergic neurons were critical for Xe-mediated improvements of autism-like behaviors. **(A)** Representative images showing CaMKII (green) and c-Fos (red) co-localization in the PVT following xenon (Xe) exposure. **(B)** Quantification of CaMKII^+^ neurons co-expressing c-Fos in the PVT. **(C)** Experimental timeline for viral injection and fiber photometry recordings of calcium activity. **(D)** Quantification of calcium activity. **(E)** Time-course of calcium activity during social interaction. **(F)** Heatmap illustrating dynamic calcium activity during social interaction. **(G)** Experimental timeline for viral injection, Xe exposure, clozapine-N-oxide (CNO) administration, and behavioral testing in L35P^+/−^ mice. **(H–J)** Representative movement traces or heatmaps (left) and quantification (right) of the discrimination index in the novel object recognition test (NORT) **(H)**, social novelty index in the three-chamber social test (TCST) **(I)**, and novel arm preference index in the Y-maze test (YMT) **(J)**. **(K)** Experimental timeline for viral injection, CNO administration, and behavioral testing in L35P^+/−^ mice. **(L–N)** Representative movement traces or heatmaps (left) and quantification (right) of the discrimination index in NORT **(L)**, social novelty index in TCST **(M)**, and novel arm preference index in YMT **(N)**. Data are presented as mean ± SEM. \**P* < 0.05, \*\**P* < 0.01, \*\*\**P* < 0.001 and \*\*\*\**P*; ^#^*P* < 0.05, ^##^*P* < 0.01, ^###^*P* < 0.001 and ^####^*P* < 0.001.

Next, we monitored the activity of PVT glutamatergic neurons during social interactions using fiber photometry. An AAV-CaMKII-GCaMP6 virus was injected into the PVT, and neuronal activity was recorded following a 21-day recovery period. Compared with WT mice, L35P^+/−^ mice exhibited reduced PVT glutamatergic neuronal activity during social interaction (−0.49 ± 0.04 vs. 0.45 ± 0.08 in controls, p < 0.0001; Fig. 2D-F). Of note, Xe exposure restored this reduced PVT neuronal activity in the L35P^+/−^ mice during social interaction (0.27 ± 0.04 vs. −0.49 ± 0.04 in controls, p < 0.0001; Fig. 2D-F).

To test whether PVT glutamatergic neuronal activity is required for Xe-mediated behavioral improvements, L35P^+/−^ mice were injected with AAV-CaMKII-hM4D(Gi) to chemogenetically inhibit PVT glutamatergic neurons during Xe exposure (Fig. 2G). After a 15-day recovery period, mice were treated with clozapine-N-oxide (CNO; 1 mg/kg, i.p.) 30 min before daily Xe exposure for 10 consecutive days, followed by behavioral assessments.

Inhibition of PVT glutamatergic neurons abolished Xe-induced improvements in discrimination index in the NORT and social novelty index in the TCST, while no significant changes was observed in the novel arm preference index in the YMT compared with control mice (Fig. 2H-J). Conversely, chemogenetic activation of PVT glutamatergic neurons mimicked the therapeutic effects of Xe in L35P^+/−^ mice (Fig. 2K). This activation also increased the discrimination index in the NORT (0.62 ± 0.04 vs. 0.45 ± 0.04 in controls, p < 0.05; Fig. 2L), social novelty index in the TCST (0.61 ± 0.03 vs. 0.53 ± 0.01 in controls, p < 0.05; Fig. 2M), and novel arm preference index in the YMT (0.60 ± 0.04 vs. 0.51 ± 0.02 in controls, p < 0.05; Fig. 2N) compared with control mice.

### Xe reduced GABA transmission through inhibition of glutamic acid decarboxylase expression in PVT inhibitory neurons

L35P^+/−^ mice were exposed to Xe or control air for 10 min per day over 10 consecutive days, after which PVT regions were dissected for single-cell RNA sequencing (Fig. 3A). Cell-type annotation showed that the PVT comprised multiple cell population including excitatory neurons, inhibitory neurons, cholinergic neurons, astrocytes, microglia, oligodendrocytes and endothelial cells. Among these, excitatory glutamatergic neurons were the most abundant, accounting for approximately 50% of all cells (Fig. 3B-C). Xe exposure induced distinct differential gene expression (DGE) profiles across multiple cell types, with particularly prominent changes observed in inhibitory neurons (Fig. 3D–E). Functional enrichment analysis showed that DGE in inhibitory neurons was associated with synapse-related pathways, including translation at the pre-synapse, synapse, and post-synapse, as well as glutamate receptor activity (Fig. 3F–G). Of note, Xe treatment reduced the expression of *glutamic acid decarboxylase 1* (*Gad1*) and *glutamic acid decarboxylase 2* (*Gad2*) in PVT inhibitory neurons; this was further validated using qRT-PCR (Fig. 3I).

**Fig. 3.**
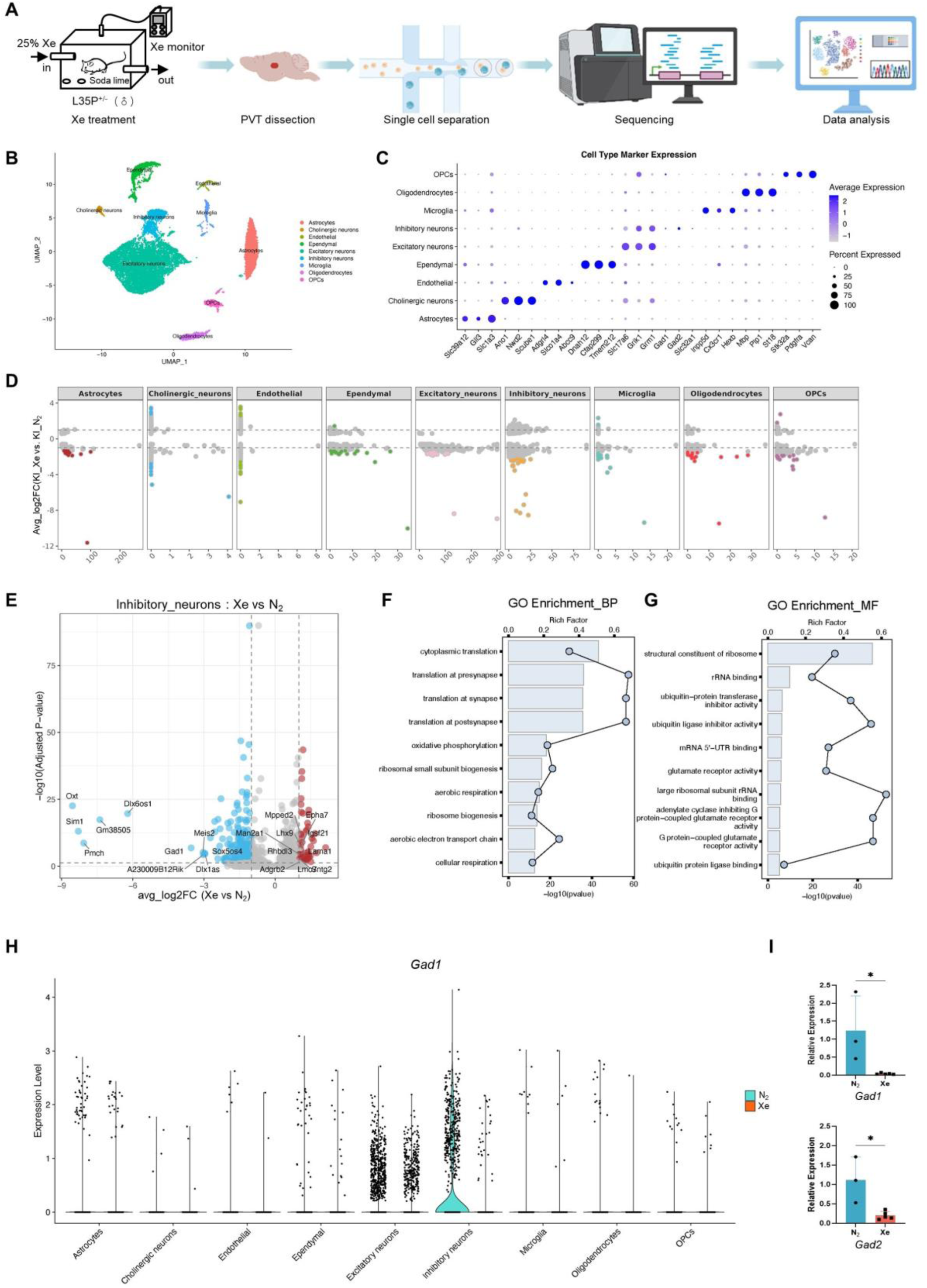
Xe decreased *Gad* expression in PVT inhibitory neurons of L35P^+/-^ mice. **(A)** Schematic of xenon (Xe) exposure and single-cell RNA sequencing performed after treatment. **(B)** Clustering of PVT cell types in L35P^+/-^ mice exposed to nitrogen (N_2_). (**C**) Representative plot showing cell-type identification based on canonical marker genes. **(D)** Differentially expressed genes (DEGs) across different PVT cell types between groups. **(E)** DEGs identified in inhibitory neurons between the two groups. **(F–G)** Gene Ontology (GO) enrichment analysis of DEGs in the PVT inhibitory neurons, including biological processes (BP) **(F)** and molecular functions (MF) **(G)**. **(H)** Expression levels of *Gad1* across different PVT cell types in both groups. **(I)** qRT–PCR analysis of *Gad1* and *Gad2* expression in both groups. Data are presented as mean ± SEM. \**P* < 0.05.

Furthermore, GABA transmission onto glutamatergic neurons was investigated using whole-cell patch-clamp recordings. L35P^+/−^ mice treated with Xe showed an increased frequency of action potentials (APs) (Fig. 4A-B), along with a decreased resting membrane potential (−64.21 ± 3.98 vs. −67.80 ± 3.38 in control, p < 0.05, Fig. 4C) and lower rheobase (36.43 ± 15.98 vs. 59.17 ± 18.32 in control, p < 0.05, Fig. 4D) in PVT glutamatergic neurons compared to controls. In addition, Xe decreased the frequency of miniature inhibitory postsynaptic currents (mIPSCs) (1.47 ± 0.41 pA vs. 2.71 ± 1.00 pA in control, p < 0.05, Fig. 4E-F) and shifted the cumulative probability distribution of inter-event intervals (Fig. 4G), without affecting mIPSCs amplitude or the cumulative probability distribution of amplitudes (Fig. 4H-I).

**Fig. 4.**
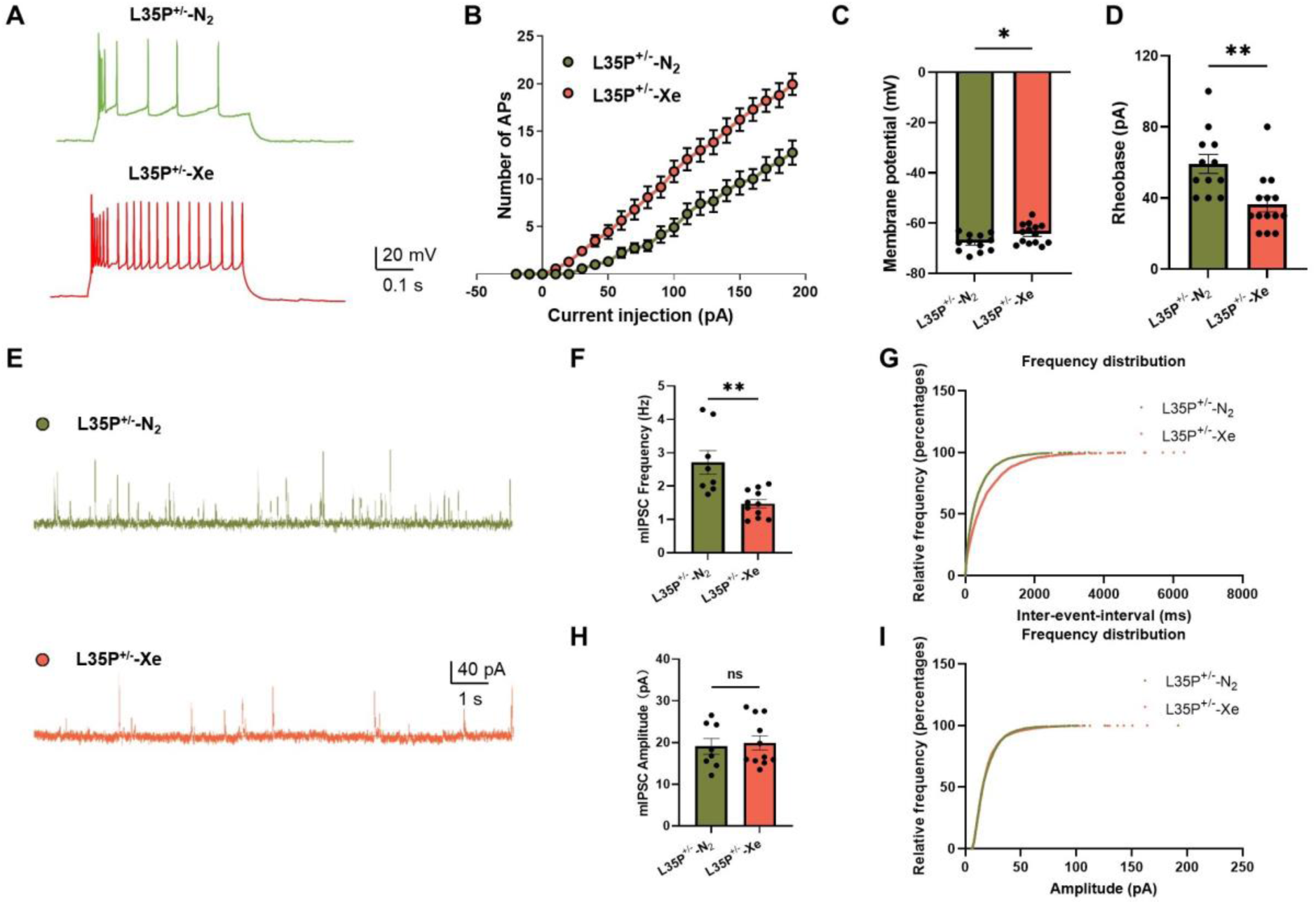
Xe increased PVT glutamatergic neuronal activity in L35P^+/-^ mice. **(A)** Presentative traces of APs evoked by depolarizing currents in L35P^+/-^ mice exposed to nitrogen (N_2_) or xenon (Xe). **(B)** Number of APs at each depolarizing current step in PVT glutamatergic neurons. **(C-D)** Quantification of the resting membrane potential amplitude **(C)** and rheobase **(D)** in both groups. **(E)** Representative traces of mIPSCs of PVT glutamatergic neurons in both groups. **(F-I)** Quantification of mIPSCs frequency **(F)**, cumulative probability distribution of inter-event intervals **(G)**, mIPSCs amplitude **(H)** and cumulative probability distribution of amplitudes **(I)** in both groups. AP, action potential; mIPSCs, miniature inhibitory postsynaptic currents. Data are presented as mean ± SEM. \**P* < 0.05, and \*\**P* < 0.01.

### GABA transmission contributed to Xe-mediated improvements in autism-like behaviors

The *Gad1* gene encodes glutamic acid decarboxylase 1 (Gad1), a key enzyme responsible for the conversion of glutamate into γ-aminobutyric acid (GABA). To examine PVT GABA release dynamics during social interaction, fiber photometry recordings were conducted (Fig. 5A).

**Fig. 5.**
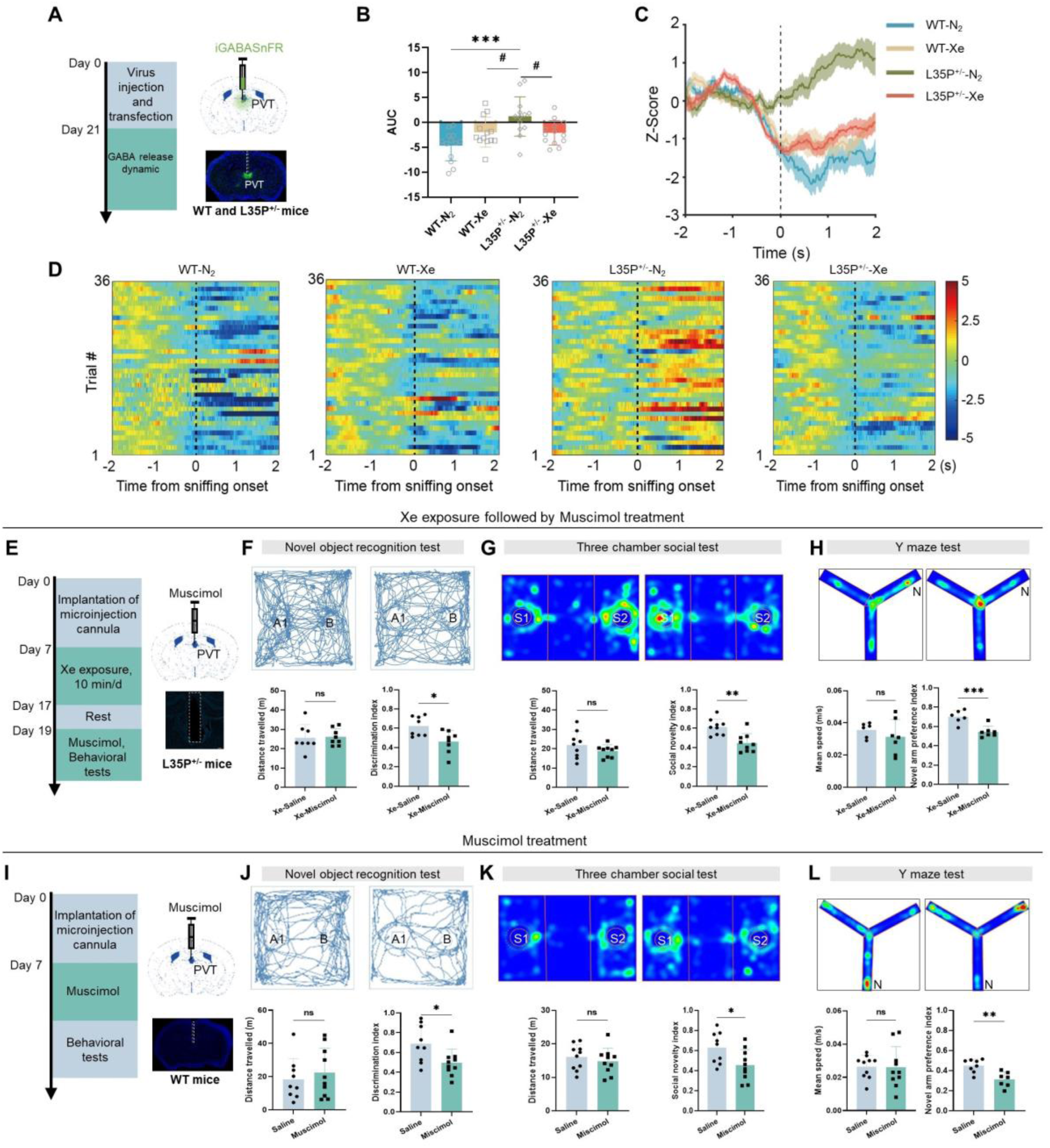
GABA transmission contributed to Xe-mediated improvements of autism-like behaviors. **(A)** Experimental timeline for viral injection and fiber photometry recordings of GABA release during social interaction in wild-type (WT) mice and L35P^+/-^ mice either exposed to N_2_ or Xe. **(B)** Quantification of GABA release. **(C)** Time-course of GABA release. **(D)** Heatmap illustrating dynamic GABA release. **(E)** Experimental timeline for Xe exposure followed by Muscimol mirco-infusion, and behavioral testing in WT mice. **(F–H)** Representative movement traces or heatmaps (left) and quantification (right) of the discrimination index in the novel object recognition test (NORT) **(F)**, social novelty index in the three-chamber social test (TCST) **(G)**, and novel arm preference index in the Y-maze test (YMT) **(H)**. **(I)** Experimental timeline for Muscimol mirco-infusion, and behavioral testing in WT mice. **(J–L)** Representative movement traces or heatmaps (left) and quantification (right) of the discrimination index in the novel object recognition test (NORT) **(J)**, social novelty index in the three-chamber social test (TCST) **(K)**, and novel arm preference index in the Y-maze test (YMT) **(L)**. Data are presented as mean ± SEM. \**P* < 0.05, and \*\*\*\**P*; ^####^*P* < 0.001.

GABA release during social interaction was increased in L35P^+/−^ compared with WT mice (Fig. 5B-D). Of note, Xe exposure normalized this elevated GABA release (0.040 ± 0.01 vs. 1.24 ± 0.56 in controls, p < 0.05), restoring it to the levels comparable to those observed in control animals (Fig. 5B-D).

To determine whether GABAergic transmission regulated Xe-mediated improvements in autism-like behaviors, L35P^+/−^ mice were treated with the GABA receptor agonist Muscimol, followed by Xe exposure (Fig. 5E). Activation of PVT GABA receptor signaling abolished Xe-induced improvements in the discrimination index in the NORT (0.46 ± 0.11 vs. 0.62 ± 0.10, Fig. 5F), the social novelty index in the TCST (0.45 ± 0.10 vs. 0.61 ± 0.08, Fig. 5G) and the novel arm preference index in the YMT (0.54 ± 0.06 vs. 0.70 ± 0.07, Fig. 5H) when compared with control mice. Conversely, activation of PVT GABA receptor in WT mice induced autism-like behaviors that mimicked those observed in L35P^+/−^ mice, as evidenced by decreased discrimination index in the NORT (0.50 ± 0.04 vs. 0.69 ± 0.06 of the controls, p < 0.05, Fig. 5J), decreased social novelty index in the TCST (0.46 ± 0.06 vs. 0.64 ± 0.04 of the controls, p < 0.05, Fig. 5K) and decreased novel arm preference index in the YMT related to controls (0.24 ± 0.04 vs. 0.41 ± 0.03, p < 0.01, Fig. 5L).

### PVT-CeA circuit drove Xe-mediated improvements in autism-like behaviors

PVT glutamatergic neurons innervate multiple downstream brain regions and play critical roles in memory and social information processing(*7, 9*). To map the projection patterns of PVT glutamatergic neurons, an AAV-CaMKII-eGFP virus was injected into the PVT (Fig. 6A-B). We found widespread projections of PVT glutamatergic neurons to multiple brain regions, including high-density projections to the zona incerta (ZI), moderate-density projections to the central amygdala (CeA), medial part of the anterior olfactory nucleus, (AOM), medial orbital cortex (MO), and core of the nucleus accumbens (AcbC). In addition, there were low-density projections to the stria terminalis (ST), anterior part of the basolateral amygdaloid nucleus (BLA), and the prelimbic cortex (PrL) (Fig. 6C-D).

**Fig. 6.**
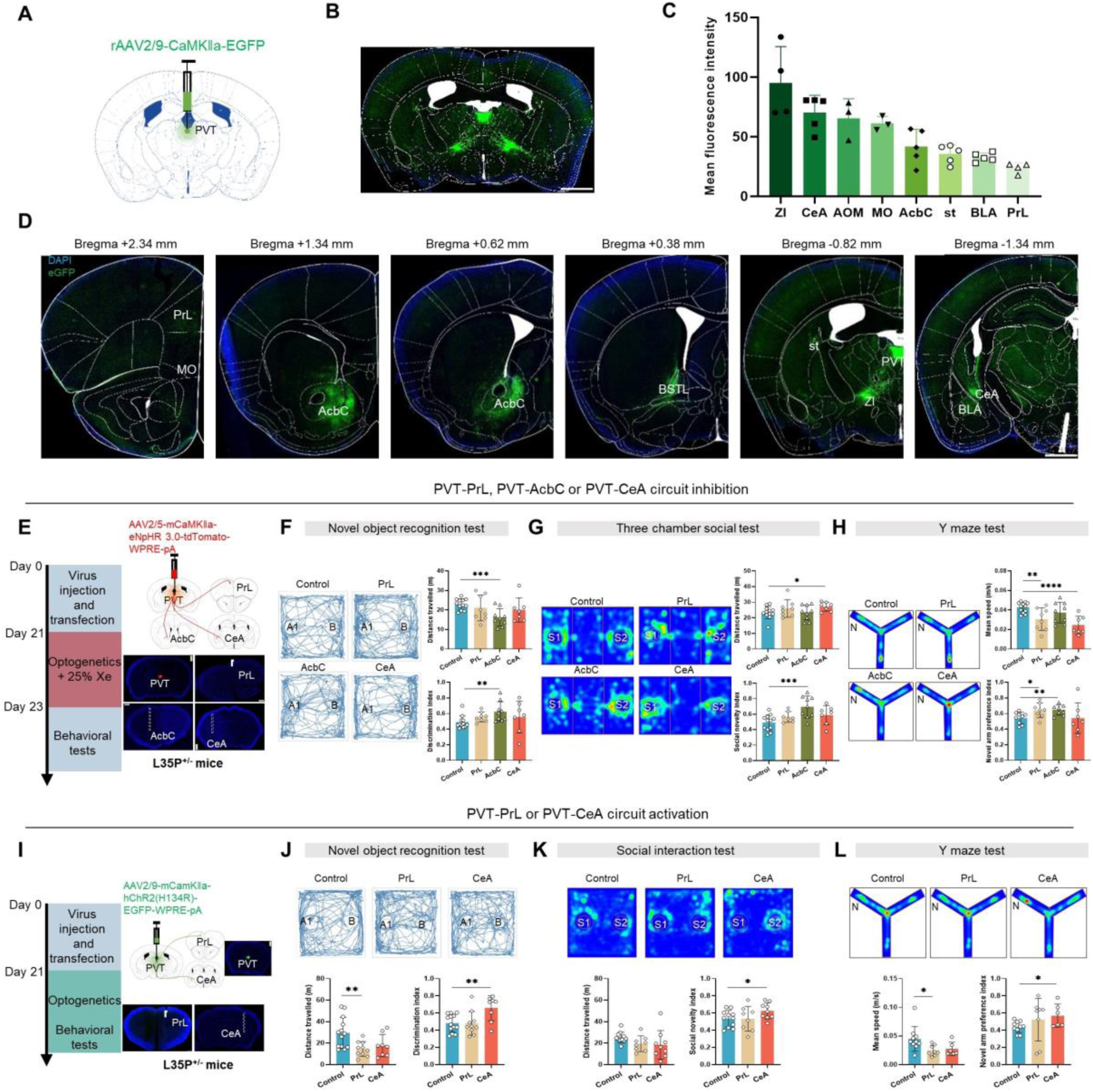
PVT-CeA circuit gated Xe-mediated improvements of autism-like behaviors. **(A)** AAV2/9-CaMKII-EGFP was injected into PVT of wild-type (WT) mice. **(B)** Representative image showing virus transfection. **(C)** Quantification of mean fluorescence intensity of the downstream regions projected by PVT glutamatergic neurons. **(D)** Representative images showing the projected downstream regions of the PVT glutamatergic neurons. ZI, zona incerta; CeA, central amygdala; AOM, anterior olfactory nucleus, medial part; MO, medial orbital cortex; AcbC, nucleus accumbens; st, stria terminalis; BLA, basolateral amygdaloid nucleus, anterior part; PrL, prelimbic cortex. **(E)** Experimental timeline for viral injection, Xe exposure, optogenetics, and behavioral testing in L35P^+/−^ mice. **(F–H)** Representative movement traces or heatmaps (left) and quantification (right) of the discrimination index in the novel object recognition test (NORT) **(F)**, social novelty index in the three-chamber social test (TCST) **(G)**, and novel arm preference index in the Y-maze test (YMT) **(H)**. **(I)** Experimental timeline for viral injection, optogenetics, and behavioral testing in L35P^+/−^ mice. **(J–L)** Representative movement traces or heatmaps (left) and quantification (right) of the discrimination index in NORT **(J)**, social novelty index in TCST **(K)**, and novel arm preference index in YMT **(L)**. Data are presented as mean ± SEM. \**P* < 0.05, \*\**P* < 0.01 and \*\*\*\**P*< 0.0001.

**Fig. 7.**
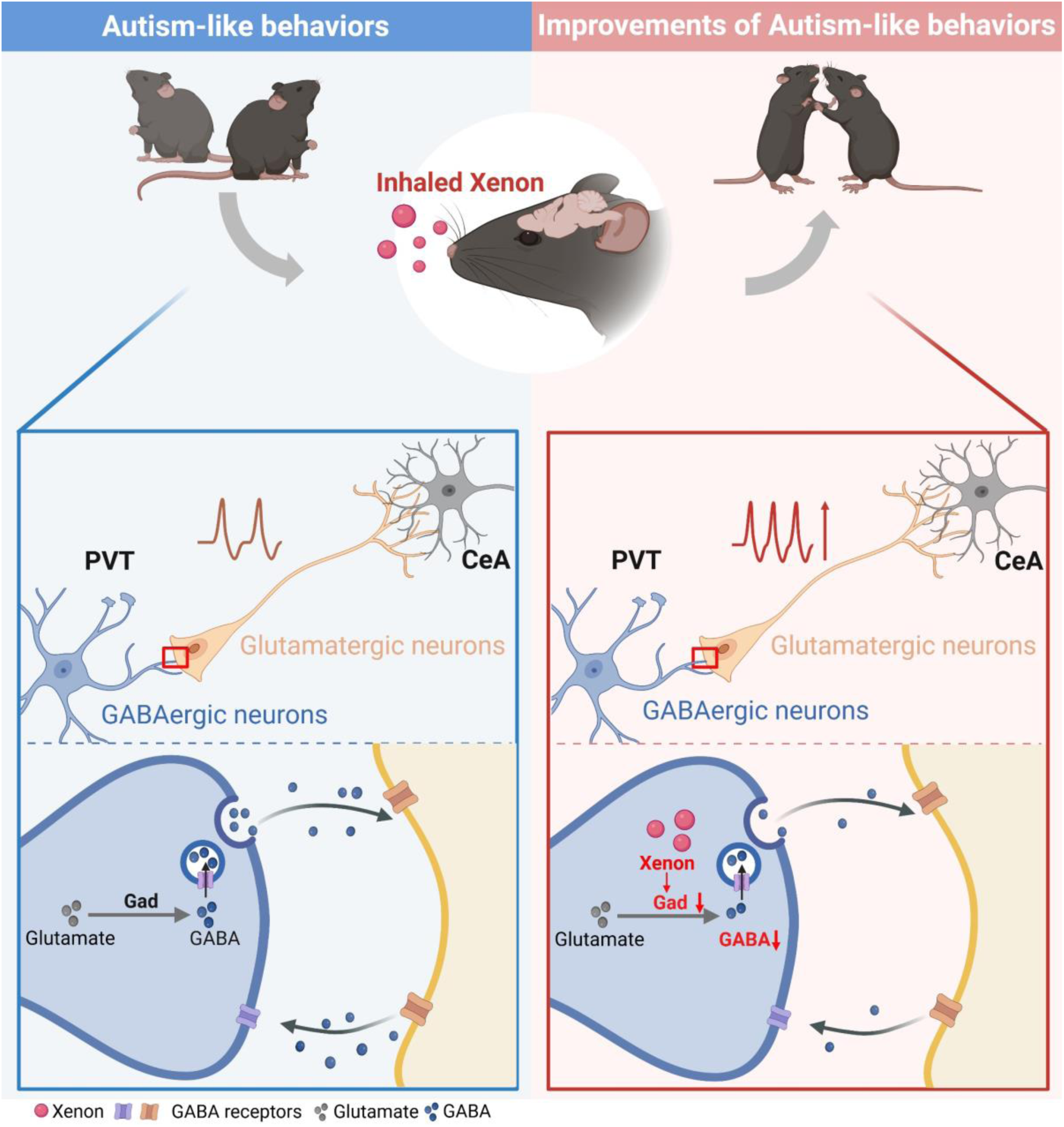
Xenon improves autism-like behaviors in mice through potentiation of PVT–CeA glutamatergic neurotransmission. In ASD mouse models, *Gad* expression was increased in inhibitory neurons of the paraventricular thalamic nucleus (PVT), leading to increased GABA release and suppression of glutamatergic neuronal activity projecting from the PVT to the central amygdala (CeA), thereby contributing to ASD-like behaviors. Xe treatment decreased Gad expression in PVT inhibitory neurons and reduced GABA release, which enhanced the activity of PVT glutamatergic neurons projecting to the CeA and ultimately improved ASD-like behaviors. Xe, xenon; ASD, autism spectrum disorder; Gad, glutamic acid decarboxylase; PVT, paraventricular thalamic nucleus; CeA, central amygdala; GABA, γ-aminobutyric acid.

To determine which downstream PVT glutamatergic projections are involved in Xe-mediated improvements in autism-like behaviors, we selected representative downstream targets with high (CeA), moderate (AcbC), and low (PrL) projection densities for circuit-specific manipulation. An AAV-CaMKII-eNpHR virus was injected into the PVT of in L35P^+/−^ mice and optical fibers were implanted above the PrL, AcbC, or CeA (Fig. 6E). After a 21-day recovery period, mice were exposed to Xe daily for 10 consecutive days, during which optogenetic inhibition of specific PVT projections was applied, followed by behavioral testing (Fig. 6E). When compared to control, optogenetic inhibition of the PVT–PrL glutamatergic projection during Xe exposure abolished Xe-induced improvements in the discrimination index in the NORT (Fig. 6F) and the social novelty index in the TCST (Fig. 6G); there were no significant improvements in the novel arm preference index in the YMT (Fig. 6H). Inhibition of the PVT–AcbC glutamatergic projection during Xe exposure did not block xenon-induced improvements in autism-like behaviors (Fig. 6F-H). Strikingly, optogenetic inhibition of the PVT–CeA glutamatergic projection during Xe exposure abolished Xe-induced improvements in autism-like behaviors, as shown by increased discrimination index in the NORT (0.56 ± 0.08 vs. 0.49 ± 0.02 in controls, p>0.05; Fig. 6F), social novelty index in the TCST (0.59 ± 0.05 vs. 0.49 ± 0.03 in controls, p>0.05; Fig. 6G), and novel arm preference index in the YMT (0.54 ± 0.07 vs. 0.54 ± 0.03 in controls, p>0.05; Fig. 6H) when compared with control mice.

An AAV-CaMKII-hChR2 virus was injected into the PVT of in L35P^+/−^ mice and optical fibers were implanted above the Pr or CeA, then followed by behavioral tests (Fig. 6I). Optogenetic activation of the PVT–PrL projection did not mimic Xe-mediated behavioral improvements as no significant changes were observed in NORT, TCST, or YMT performance (Fig. 6I-L).

Conversely, optogenetic activation of the PVT–CeA circuit was sufficient to mimic Xe-mediated therapeutic effects, resulting in improvements in NORT, TCST, and YMT performance (Fig. 6I-L). Finally, c-Fos immunostaining showed that Xe exposure increased neuronal activity in the CeA (fig. S6A-B). Double-labeling analysis further demonstrated that Xe-activated neurons in the CeA were predominantly GABAergic (fig. S6C-D), suggesting that Xe improves autism-like behaviors through a PVT^Glu^–CeA^GABA^ circuit.

## DISCUSSION

In this study, we probed the molecular, cellular and circuit-level mechanisms by which Xe ameliorates autism spectrum disorder (ASD)–related phenotypes across multiple ASD mouse models. We demonstrated that Xe reduced the expression of glutamate decarboxylase (Gad), the enzyme responsible for converting glutamate to γ-aminobutyric acid (GABA), leading to decreased GABA release from paraventricular thalamus (PVT) GABAergic neurons and a corresponding enhancement of PVT glutamatergic neuronal activity. Of note, Xe-enhanced glutamatergic transmission from the PVT to inhibitory neurons in the central amygdala (CeA) and rescued deficits in ASD-associated behaviors. These findings support Xe as a promising and translatable therapy for ASD.

Previous studies have demonstrated the neuroprotective potential of Xe. In a proof-of-concept, open-label, randomized controlled trial, Xe administration was shown to be both feasible and apparently safe after birth asphyxia(*23*). In addition, inhaled Xe combined with therapeutic hypothermia, compared with hypothermia alone, resulted in reduced white matter damage in comatose survivors (24-27 years of age) of out-of-hospital cardiac arrest (*24*). Consistent with these findings, our previous open-label study demonstrated that Xe represents a potentially effective acute treatment modality for panic disorder(*24*). Moreover, in a preclinical valproic acid (VPA)–induced rodent model of autism, we showed that acute inhalation of 25% Xe for 10 min improved social behavior, reduced exploratory motivation, and normalized performance in the forced swim test(*21*). However, our preliminary report did not explore the underlying mechanisms. In the currently study, we have clearly demonstrated that acute inhalation of 25% Xe in genetic mouse models of ASD ameliorated ASD-like behaviors, particularly learning and memory deficits. Of note, repeated Xe inhalation at the same concentration (10 min per day for 10 consecutive days) produced sustained behavioral improvements that persisted for at least 10 days after cessation of exposure, suggesting that long-term Xe administration may represent a viable therapeutic strategy for ASD.

Although the precise mechanisms by which Xe modulates neuronal activity remain incompletely understood, accumulating evidence suggests that Xe plays an important role in synaptic transmission, particularly in glutamatergic and GABAergic signaling. Previous studies showed that acute exposure to 60% Xe selectively inhibits the N-Methyl-D-Aspartate (NMDA) receptor–mediated component of glutamatergic excitatory postsynaptic currents (*18*). Furthermore, 50% or 70% Xe inhibits α-amino-3-hydroxy-5-methyl-4-isoxazole-4-propionic acid (AMPA) receptor-mediated glutamatergic excitatory transmission in spinal lamina IX neurons via a postsynaptic mechanism(*25*). In contrast to these acute, high-dose inhibitory effects, we found that long-term administration of 25% Xe in ASD mouse models enhanced glutamatergic neuronal activity in the PVT. Importantly, inhibition of PVT glutamatergic activity abolished the behavioral improvements induced by Xe, indicating that facilitation of glutamatergic transmission is required for its therapeutic efficacy. Collectively, these findings suggest that Xe modulates glutamatergic signaling through distinct mechanisms that depend on dose, duration of exposure, and region- or cell-type–specific actions.

We further found that long-term Xe administration in mouse ASD models selectively reduced the expression of glutamate decarboxylase in GABAergic neurons, as revealed by single-cell RNA sequencing. Gene Ontology enrichment analysis further indicated that Xe-responsive pathways were significantly enriched in pre-synaptic, synaptic, and post-synaptic compartments, supporting a synapse-centered mechanism of action. GABA release was also increased during social interaction with a stranger mouse, and this activity-dependent increase was attenuated following Xe exposure. Potentiation of GABA_A_ receptor activity *via* direct agonist injection into the PVT attenuated Xe-induced behavioral improvements. This agrees with previous findings(*26*) showing that Xe reduces the frequency, but not the amplitude, of spontaneous inhibitory post-synaptic currents, along with a depolarized resting membrane potential and lower rheobase; hallmarks of a pre-synaptic mechanism of action. These findings suggest that Xe ameliorates ASD-like behaviors by selectively reducing GABA synthesis and dampening excessive GABAergic transmission, thereby restoring excitatory–inhibitory balance in ASD.

PVT glutamatergic neurons play critical roles in regulating social behavior, memory, wakefulness, and emotional processing. Using viral-based neural tracing approaches, we found that PVT glutamatergic neurons project to multiple downstream regions with varying densities, including high-density projections to the zona incerta (ZI), moderate-density projections to the central amygdala (CeA) and nucleus accumbens (AcbC), and low-density projections to the prelimbic cortex (PrL). Previous studies demonstrated that PVT-AcbC is involved in stress-related social behaviors, such that inhibition of the PVT–AcbC circuit exacerbates stress-induced social deficits, whereas activation of this pathway alleviates these impairments(*8*). However, in our mouse ASD models, inhibition of the PVT–AcbC circuit did not abolish Xe-induced behavioral improvements. A similar lack of effect was observed following inhibition of the PVT–PrL circuit. In contrast, inhibition of the PVT–CeA circuit completely abolished the beneficial behavioral effects of Xe. Moreover, direct activation of the PVT–CeA circuit in the absence of Xe was sufficient to improve ASD-like behaviors, closely mimicking the effects of Xe administration. Consistent with this finding, previous studies showed that the PVT–CeA circuit is also involved in regulating both depression-related behaviors(*27*) and stress-induced wakefulness(*28*). More than 90% of neurons in the CeA are GABAergic, and modulation of these neuronal populations plays a central role in emotional regulation(*29*). In line with this, we found that PVT glutamatergic terminals predominantly target CeA GABAergic neurons, further supporting a model in which Xe exerts its beneficial effects through modulation of the PVT^Glu^–CeA^GABA^ circuit.

In summary, we have demonstrated that inhaled xenon ameliorates ASD-like phenotypes in mouse models by modulating the activity of the PVT^Glu^–CeA^GABA^ circuit, supporting further clinical application of xenon as a potential therapy for ASD. A multicenter, randomized, controlled trial to evaluate the efficacy of Xe treatment in children with ASD is scheduled to begin in 2026 (NCT07435103).

## Author contributions

Conceptualization: JPZ, QS, DQM

Methodology: YNR, JTW, XKL

Investigation: JPZ, YNR, JTW, XX, YYL, HFL

Visualization: RYN, XX, ZAW, JBT, YWO

Funding acquisition: JPZ, QS, DQM

Supervision: JPZ, DQM

Writing – original draft: YNR, JPZ,

Writing – review & editing: JPZ, DQM

## Data availability

All data generated or analyzed during this study are included in the published article (and its supplementary information files). Source data are provided in the paper and are available upon reasonable request to the corresponding authors. Single-cell sequencing data are available https://ngdc.cncb.ac.cn/gsa/s/IOReK2Pd.

## SUPPLEMENTARY MATERIALS

Materials and Methods Figs. S1 to S6

