## Supplementary Materials for "Xenon improves autism-like behaviors in mice through potentiation of PVT–CeA glutamatergic neurotransmission"

### **MATERIALS AND METHODS**

#### **Animals**

All experiments were conducted in accordance with the ARRIVE Guideline and the experimental protocol was approved by the Ethics Committee of the Laboratory Animal Center of Zhejiang University (NO. ZJU20241110). Eight-week-old genetic models including *Mef2c* mutant (L35P<sup>+/-</sup>) mice and *Shank3* knockout (*Shank3*<sup>-/-</sup>) mice were obtained from Shulaibao Biotechnology (Wuhan, China), and C57BL/6J mice were purchased from Changsheng Biotechnology (Liaoning, China). Five mice per cage (same sex) were housed under specific pathogen-free conditions (temperature 23 ± 1 °C, humidity 50 ± 5%) with a 12-h light/dark cycle and free access to food and water.

#### **Xenon inhalation**

A purposed built Xe inhalational system was constructed consisting of a sealed chamber capable of accommodating up to six mice per treatment, connected to an inlet tube for gas delivery and an outlet tube for gas release. The bottom of the chamber was filled with sodium hydroxide to absorb carbon dioxide. Xe (NO. 7440-63-3, NEWRADAR, Wuhan, China) concentration inside the chamber was continuously monitored using a gas monitor (NO. MS400, JIENSI, Guangzhou, China). Mice were randomly assigned to two groups including the Xe exposure group (Xe) and the control group (N<sub>2</sub>). Mice in the Xe group were exposed to a gas mixture containing 25% Xe and 30% oxygen (O<sub>2</sub>) balanced with nitrogen (N<sub>2</sub>) for 10 min per day over 10 consecutive days. Mice in the N<sub>2</sub> group were exposed to a gas mixture containing 0% Xe, 30% O<sub>2</sub>, and 70% N<sub>2</sub> for 10 min per day over the same period. For short-term exposure experiments, mice in the Xe group received a single 30-min exposure to 25% Xe + 30% O<sub>2</sub> + 45% N<sub>2</sub>, whereas control mice were exposed to 0% Xe + 30% O<sub>2</sub> + 70% N<sub>2</sub> for 30 min.

#### **Behavioral tests**

Behavioral tests including the novel object recognition test (NORT), three-chamber social test (TCST), and Y-maze test (YMT) were conducted between 9 am. and 5 pm. Mice were handled for one week before experiments and were transported to the behavioral testing room at least 2 hours prior to assessment to minimize stress. The behavioral apparatus was thoroughly cleaned with 75% ethanol between trials. During each behavioral assessment, mice were allowed to move freely and explore the apparatus. Their behaviors were recorded and analyzed offline using the Any-maze video-tracking system (Stoelting Co., Wood Dale, Illinois, USA).

##### *Novel object recognition test*

The novel object recognition test (NORT) was conducted in an open-field apparatus (40 × 40 × 40 cm). During the habituation stage, two identical objects were placed on opposite sides of the chamber, and the mouse was placed in the center zone and allowed to freely explore both objects for 10 min. After the habituation session, the mouse was returned to its home cage for a 1-h retention interval. In the testing stage, one of the familiar objects was replaced with a novel object, and the mouse was again placed in the center of the chamber and allowed to explore the two objects for 10 min. The discrimination index (DI) was calculated as the ratio of the time spent exploring the novel object to the total exploration time of both objects: $DI = T_n / (T_n + T_f)$, where $T_n$ represents the time spent exploring the novel object and $T_f$ represents the time spent exploring the familiar object.

##### *Three-chamber social test*

The TCST was conducted in a behavioral apparatus (60 × 40 × 30 cm) consisting of three equal-sized compartments separated by opaque dividers. During the habituation session, both end chambers were closed by the dividers, and the test mouse was placed in the center chamber and allowed to habituate for 5 min. For the sociability test, an empty wire cup and a wire cup containing an unfamiliar age- and sex-matched mouse (Stranger I) were placed in the left and right chambers, respectively. The dividers were then removed, allowing the test mouse to freely explore all three chambers for 10 min. To assess social novelty preference, a second unfamiliar mouse (Stranger II) was placed under the previously empty wire cup, while Stranger I remained in the original location. The test mouse was again allowed to explore all three chambers for 10 min, and its behavior was recorded. The social novelty index was calculated as $T_2 / (T_1 + T_2)$, where $T_2$ represents the time spent interacting with Stranger II and $T_1$ represents the total time spent interacting with Stranger I and Stranger II.

#### *Y-maze test*

The Y-maze test (YMT) was conducted in a behavioral apparatus consisting of three arms. One arm was randomly designated as the novel arm and was initially blocked by a barrier. During the training phase, the mouse was allowed to freely explore the other two arms for 10 min. In the testing phase, 1 hour after training, the barrier was removed to allow access to the previously blocked novel arm, and the mouse was allowed to explore all three arms for 5 min. The novel arm preference index was calculated as $N/(N + F)$, where N represents the time spent in novel arm and F represents the time spent in the two previous familiar arms.

#### **Stereotaxic surgery**

Mice were anesthetized with 2% isoflurane (RWD Life Science, Shenzhen, China) and secured in a digital stereotaxic apparatus (68018, RWD Life Science, Shenzhen, China) for viral injection. For the paraventricular thalamus (PVT), the coordinates were anterior–posterior (AP) 0 mm, medial–lateral (ML) –0.94 mm, and dorsal–ventral (DV) –3.0 mm; for the central amygdala (CeA), the coordinates were AP +2.5 mm, ML –1.46 mm, and DV –4.5 mm. Micro-glass pipettes connected to a microsyringe pump (R-480, RWD Life Science, Shenzhen, China) were used to deliver AAV at a rate of 20 nL/min, with a total injection volume of 200 nL per site. After injection, the pipette was kept in place for 10 min. Animals were subsequently placed on a warming pad until they recovered fully. Mice with off-target viral expression confirmed by *post hoc* histological verification were excluded from further studies.

#### **Chemogenetics**

For chemogenetic inhibition, AAV2/9-CaMKIIα-hM4Di (NO. S0500-9-H20, Taitool, Shanghai, China) was injected into the PVT of L53P<sup>+/-</sup> mice. After a 15-day recovery period to allow for viral transfection, clozapine-N-oxide (CNO; 1 mg/kg; NO. A3317, APExBIO) was administered intraperitoneally (i.p.) 30 min before Xe exposure once daily for 10 consecutive days. Following the treatment period, mice were subjected to a battery of behavioral tests. For chemogenetic activation, AAV2/9-CaMKIIα-hM3Dq (NO. S0484-9-H20, Taitool, Shanghai, China) was injected into the PVT. After 15 days of recovery, CNO (1 mg/kg, i.p.) was administered once daily for 10 consecutive days. Behavioral test was conducted 2 days after the final treatment

### **Optogenetics**

AAV2/9-CaMKIIα-eNpHR (NO. S0577-5-H20, Taitool, Shanghai, China) was injected into the PVT of L53P<sup>+/-</sup> mice for optogenetic inhibition, and an optical fiber (Xi'an Bogao Optoelectronic Technology Co., Ltd., Xi'an, China) was implanted above the CeA. After a 21-day recovery period, light stimulation (589 nm, 5 mW, continuous illumination; Thinker Tech Nanjing Co., Ltd., Nanjing, China) was delivered for 10 min before Xe exposure once daily for 10 consecutive days. Two days after completion of the treatment, mice were subjected to a battery of behavioral tests including NORT, TCST, and YMT.

For optogenetic activation, AAV2/9-CaMKIIα-hChR2 (NO. S0826-9-H20, Taitool, Shanghai, China) was injected into the PVT of L53P<sup>+/-</sup> mice. After a 21-day recovery period, mice underwent behavioral assessments, including the NORT, social interaction test, and YMT. Mice received optical stimulation (473 nm, 10 mW, 20 Hz, 15 ms pulse width; Thinker Tech Nanjing Co., Ltd., Nanjing, China), and behavioral responses were recorded simultaneously. For the social interaction test, mice were first habituated to an open-field chamber (40 × 40 × 40 cm) for 5 min. In the sociability phase, an empty wire cup and a wire cup containing an unfamiliar age- and sex-matched mouse (Stranger I) were placed in the left and right compartments, respectively, and the test mouse was allowed to explore for 10 min. For the social novelty preference phase, a second unfamiliar mouse (Stranger II) was introduced into the previously empty wire cup, while Stranger I remained in its original location. The test mouse was then allowed to explore for an additional 10 min.

### **Fiber-photometry**

AAV2/9-CaMKIIα-GCaMP6f (NO. S0881-9-H20, Taitool, Shanghai, China) or AAV-hSyn-iGABASnFR (NO. H15052, OBiO, Shanghai, China) was injected into the PVT, and an optical fiber was implanted above the PVT for fiber photometry recording. After viral transfection, calcium activity or GABA release was recorded during the social interaction period. Signals were sampled at 40 Hz using a fiber photometry system (Thinker Tech Nanjing Co., Ltd., Nanjing, China). Data were analyzed when mice initiated an interaction with an age-matched conspecific, including a 1 s window before and a 1 s window after the onset of interaction. Fluorescence changes were calculated as $\Delta F/F = (F - F_o)/F_o$, where $F_o$ represents the mean fluorescence signal during the 1s baseline period before social interaction, and $F$ represents the peak fluorescence signal within the 1s period following onset interaction.

#### **Micro-infusion**

For cannula-guided drug microinfusion, guide cannulas (Kedou Brain-Computer Technology Co., Ltd., Suzhou, China) were implanted into the PVT of L53P<sup>+/-</sup> or C57BL/6J mice under 2% isoflurane anesthesia. Animals were allowed to recover for 7 days after surgery. L53P<sup>+/-</sup> mice received Xe treatment for 10 consecutive days. Two days after the final xenon exposure, muscimol (a GABA<sub>A</sub> receptor agonist; 1.75 mM, NO. HY-N2313, MedChemExpress, New Jersey, USA) dissolved in 0.9% saline was infused into the PVT 30 min before behavioral testing. The infusion was delivered at a rate of 0.2 μL/min for 1 min. In addition, C57BL/6J mice without Xe exposure received PVT muscimol infusion 30 min before behavioral testing.

#### **Single-cell sequencing**

Two days after Xe exposure, PVT tissue from L53P<sup>+/-</sup> mice was rapidly dissected under sterile conditions and washed in ice-cold RPMI 1640 supplemented with 0.04% BSA. Tissue was mechanically dissociated in 1 mL nuclear lysis buffer (0.1% NP-40, 10 mM Tris-HCl, 146 mM NaCl, 1 mM CaCl<sub>2</sub>, 21 mM MgCl<sub>2</sub>, 40 U/mL RNase inhibitor). Nuclear release was confirmed by trypan blue staining. An equal volume of wash buffer (10 mM Tris-HCl, 146 mM NaCl, 1 mM CaCl<sub>2</sub>, 21 mM MgCl<sub>2</sub>, 0.01% BSA, 40 U/mL RNase inhibitor) was added, and the suspension was filtered through a 40 μm strainer. The filtrate was collected, the strainer rinsed, and the combined suspension centrifuged at 500 × g for 5 min at 4°C. Nuclei were washed once in PBS containing 1% BSA and resuspended in 100 μL PBS + 1% BSA. Nuclear integrity and concentration were assessed by trypan blue staining.

Nuclei were diluted to 700–1200 nuclei/μL and processed using the Chromium Next GEM Single Cell 3' v3.1 platform (10x Genomics) according to the manufacturer's instructions. Libraries were prepared using the Chromium Single Cell 3'/5' kit and sequenced on an Illumina NovaSeq 6000 (paired-end, 150 bp).

FASTQ files were processed with Cell Ranger (v9.0.1) and aligned to the mouse genome (CRCm39). Downstream analyses were performed in R using Seurat (v4.3.0). Nuclei were retained if they contained 200–6,000 genes, 1,000–30,000 Unique Molecular Identifiers

(UMIs), and $\leq 5\%$ mitochondrial or haemoglobin transcripts. Doublets were identified with scDbtFinder (v1.16.0) and removed. Data were log-normalized, and the top 2,000 highly variable genes were selected. Principal Component Analysis (PCA) was performed, followed by batch correction using Harmony. Clustering was conducted using a graph-based approach (Leiden algorithm), and Uniform Manifold Approximation and Projection (UMAP) was used for visualization. Differentially expressed genes were identified using the Wilcoxon rank-sum test with Bonferroni correction (adjusted $P < 0.05$, $|\log_2 \text{fold change}| > 1$). Gene Ontology and Kyoto Encyclopedia of Genes and Genomes (KEGG) enrichment analyses were performed with hypergeometric testing in R.

#### **Patch clamp**

L53P<sup>+/-</sup> mice with or without Xe exposure were used for patch-clamp recordings from the PVT after 2 days of treatment. Coronal brain slices (300 $\mu\text{m}$) containing the PVT were prepared using a vibratome and ice-cold, oxygenated (95% $\text{O}_2/5\% \text{CO}_2$) protective cutting solution containing (in mM): 235 sucrose, 1.25 $\text{NaH}_2\text{PO}_4$, 2.5 KCl, 0.5 $\text{CaCl}_2$, 7 $\text{MgCl}_2$, 10 glucose, 26 $\text{NaHCO}_3$, and 5 sodium pyruvate (pH 7.3, 310 mOsm). Slices were incubated at 32°C for 30 min in oxygenated artificial cerebrospinal fluid (aCSF; pH 7.4, 310 mOsm) containing (in mM): 126 NaCl, 2.5 KCl, 26 $\text{NaHCO}_3$, 1.25 $\text{NaH}_2\text{PO}_4$, 10 D-glucose, 1 sodium pyruvate, 1 $\text{MgCl}_2$, and 2 $\text{CaCl}_2$, and were subsequently maintained at room temperature in oxygenated aCSF until recording.

For electrophysiological recordings, slices were transferred to a recording chamber and continuously perfused with oxygenated aCSF at 3–5 mL/min. Whole-cell patch-clamp recordings were obtained from PVT neurons in either voltage-clamp or current-clamp mode. For voltage-clamp recordings, patch pipettes ($\sim 3 \text{ M}\Omega$) were filled with an internal solution containing (in mM): 125 CsMeSO<sub>3</sub>, 10 HEPES, 10 EGTA, 8 TEA-Cl, 5 4-AP, 0.4 Na-GTP, 4 Na<sub>2</sub>-ATP, 5 QX-314, 1 $\text{CaCl}_2$, and 1 $\text{MgCl}_2$ (pH 7.3–7.4, 280–290 mOsm). For current-clamp recordings, pipettes ($\sim 5 \text{ M}\Omega$) were filled with an internal solution containing (in mM): 130 K-gluconate, 10 HEPES, 5 KCl, 10 phosphocreatine disodium, 0.4 Na-GTP, 4 Na<sub>2</sub>-ATP, and 1 $\text{MgCl}_2$ (pH 7.3–7.4, 280–290 mOsm). Miniature inhibitory postsynaptic currents (mIPSCs) were recorded at a holding potential of 0 mV in aCSF containing 10 $\mu\text{M}$ CNQX (NO. HY-15066, MedChemExpress, New Jersey, USA), 50 $\mu\text{M}$ D-AP5 (NO. HY-100714A, MedChemExpress, New Jersey, USA), and

1 μM tetrodotoxin (TTX, HB1034, Hello Bio, Bristol, UK). To assess neuronal excitability, a series of hyperpolarizing and depolarizing current steps (0.5 s duration) were applied in 20 pA increments from $-20$ pA to $+200$ pA. Signals were recorded using an Axopatch 700B amplifier (Molecular Devices, USA), sampled at 10 kHz, and analyzed offline using pClamp 10.7 (Molecular Devices). All recordings were performed at $32^{\circ}\text{C}$.

#### Quantitative real-time PCR

Mice were deeply anesthetized with 2% isoflurane, and rapidly killed for dissection of the PVT regions. Total RNA was extracted using TRIzol reagent (R0016, Beyotime Biotechnology, Shanghai, China). Reverse transcription was performed using the HiScript II Q RT SuperMix (R223-01, Vazyme, Nanjing, China) according to the manufacturer's instructions, with 1,000 ng of total RNA per reaction. Quantitative real-time PCR (qRT-PCR) was conducted to determine the relative mRNA expression levels of *Gad1* and *Gad2*, normalized to *Actin*, using ChamQ Universal SYBR qPCR Master Mix (Q711-02, Vazyme, China). Amplification and data acquisition were performed on a CFX Connect Real-Time PCR Detection System (Bio-Rad Laboratories, USA). The primer sequences (mouse) were as follows: *Gad1*, forward 5'-ATGATACTTGGTGTGGCGTAG-3', reverse 5'-GACTCTTCTCTTCCAGGCTATTG-3'; *Gad2*, forward 5'-GCTTTTGGTCCTTCGGATCT-3', reverse 5'-GAACTTTTGGGCCACCTG-3'; *Actin*, forward 5'-TGTGATGGTGGGAATGGGTCAGAA-3', reverse 5'-TGTGGTGCCAGATCTTCTCCATGT-3'. The PCR program consisted of an initial denaturation at $95^{\circ}\text{C}$ for 30 s, followed by 40 cycles of amplification at $95^{\circ}\text{C}$ for 10 s and $60^{\circ}\text{C}$ for 30 s. Relative mRNA expression levels for *Gad1* and *Gad2* were calculated using the $2^{-\Delta\Delta\text{CT}}$ method with *Actin* as an internal control.

#### Immunohistochemistry

Mice were deeply anesthetized following experiments then perfused transcardially with phosphate-buffered saline (PBS), followed by 4% paraformaldehyde (PFA; Merck, Darmstadt, Germany). Their brains were dissected and post-fixed in 4% PFA, followed by dehydration in 30% sucrose solution. Brain tissues were sectioned into 30 $\mu\text{m}$ coronal slices using a cryostat microtome (NX70, Thermo Fisher Scientific, Waltham, MA, USA). Brain sections were washed three times with PBS and incubated in 10% normal goat serum (SL038, Solarbio Science and Technology, Beijing, China) at room temperature to block nonspecific binding. After blocking, sections were incubated overnight at $4^{\circ}\text{C}$ with the following primary antibodies: guinea pig

anti-c-Fos (1:1,000, 226308, Synaptic Systems, Germany), rabbit anti-CaMKIIα (1:500, ab5683, Abcam, USA), rabbit anti-GABA (1:500, A2052, Sigma-Aldrich, Germany), and mouse anti-GAD67 (1:1,000, MAB5406, Sigma-Aldrich, Germany). After three washes with PBS, sections were incubated for 2 hours at room temperature with secondary antibodies including Alexa Fluor<sup>®</sup> 488–conjugated AffiniPure<sup>®</sup> Goat Anti-Rabbit IgG (H+L) (1:400, 111-545-144, Jackson ImmunoResearch, UK) and Alexa Fluor<sup>®</sup> 647–conjugated AffiniPure<sup>®</sup> Donkey Anti-Guinea Pig IgG (H+L) (1:400, 706-605-148, Jackson ImmunoResearch, UK). Sections were then mounted with anti-fade mounting medium containing 4',6-diamidino-2-phenylindole (DAPI, P0131, Beyotime Biotechnology, Shanghai, China). Images were acquired using an Olympus VS120 Virtual Slide System (Olympus, Tokyo, Japan) and a TCS SP8 confocal microscope (Leica, Wetzlar, Germany).

#### **Statistical analysis**

Data are presented as the mean ± standard error of the mean (SEM) as they were normally distributed assessed with the Shapiro–Wilk test. For two group data, either a two-tailed paired or unpaired Student's t-test was applied as appropriate. For repeated-measures data, two-way ANOVA was performed followed by Fisher's LSD or Šidák's multiple-comparisons test with Bonferroni corrections. Statistical analyses were performed using GraphPad Prism (version 9.0; GraphPad Software, USA). A statistical significance was defined as p value less than 0.05.

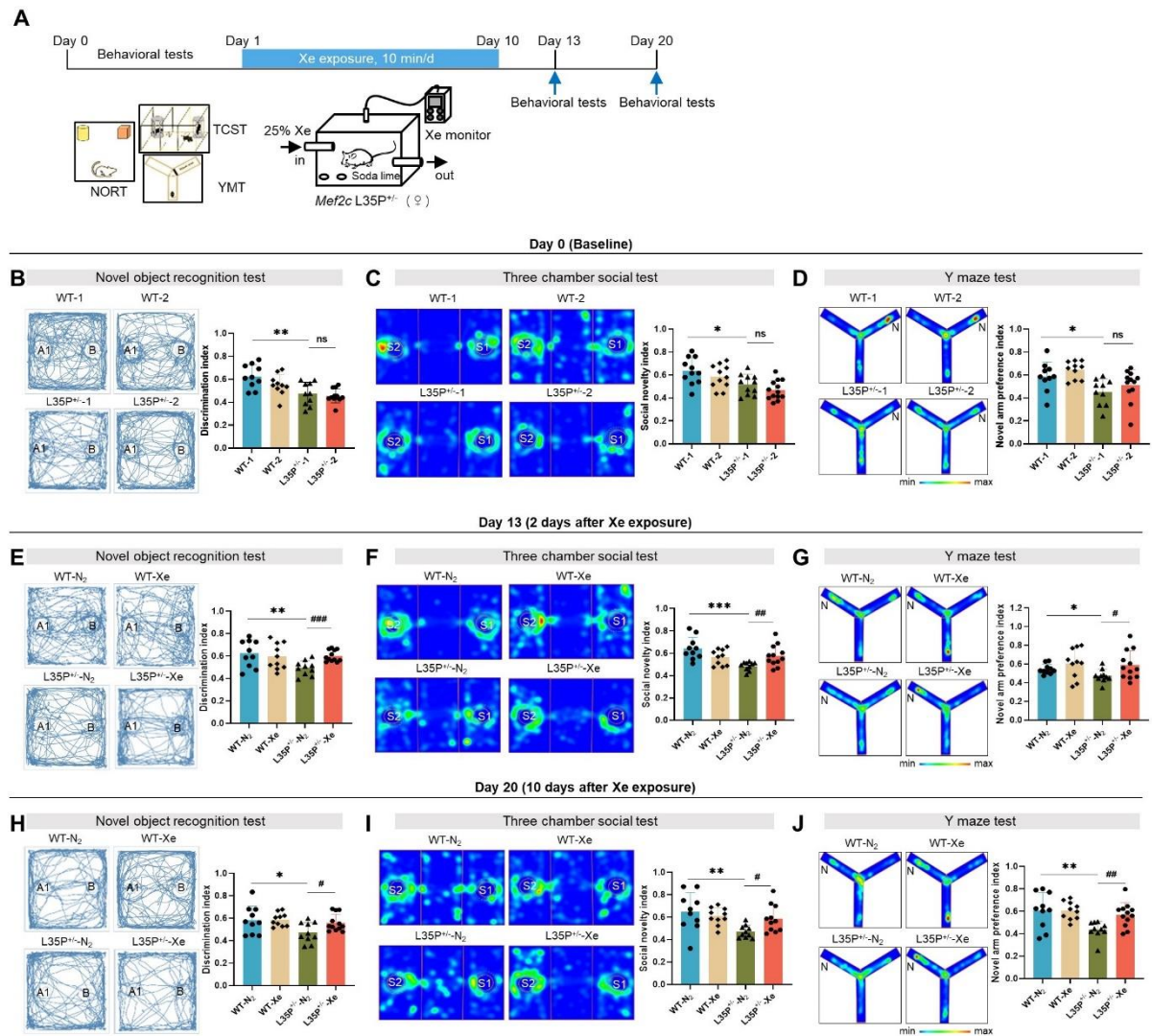

**fig. S1. Xe improved autism-like behaviors in L35P<sup>+/-</sup> female mice.** (A) Experiment timeline of xenon (Xe) exposure and behavioral tests including novel object recognition test (NORT), three-chamber social test (TCST) and Y-maze test (YMT). (B–D) Baseline behavioral performance without Xe exposure. Representative movement traces or heatmaps (left) and quantification of discrimination index in NORT (B), social novelty index in TCST (C), and novel arm preference index in YMT (D). (E–G) Behavioral performance 2 days after Xe exposure. Representative movement traces or heatmaps (left) and quantification of discrimination index in NORT (E), social novelty index in TCST (F), and novel arm preference index in YMT (G). (H–J) Behavioral performance 10 days after Xe exposure. Representative traces or heatmaps (left) and quantification of discrimination index in NORT (H), social novelty index in TCST (I), and novel arm preference index in YMT (J). Data are presented as mean ± SEM. \**P* < 0.05, \*\**P* <

0.01, and \*\*\*$P < 0.001$; #$P < 0.05$, ##$P < 0.01$, and ###$P < 0.001$.

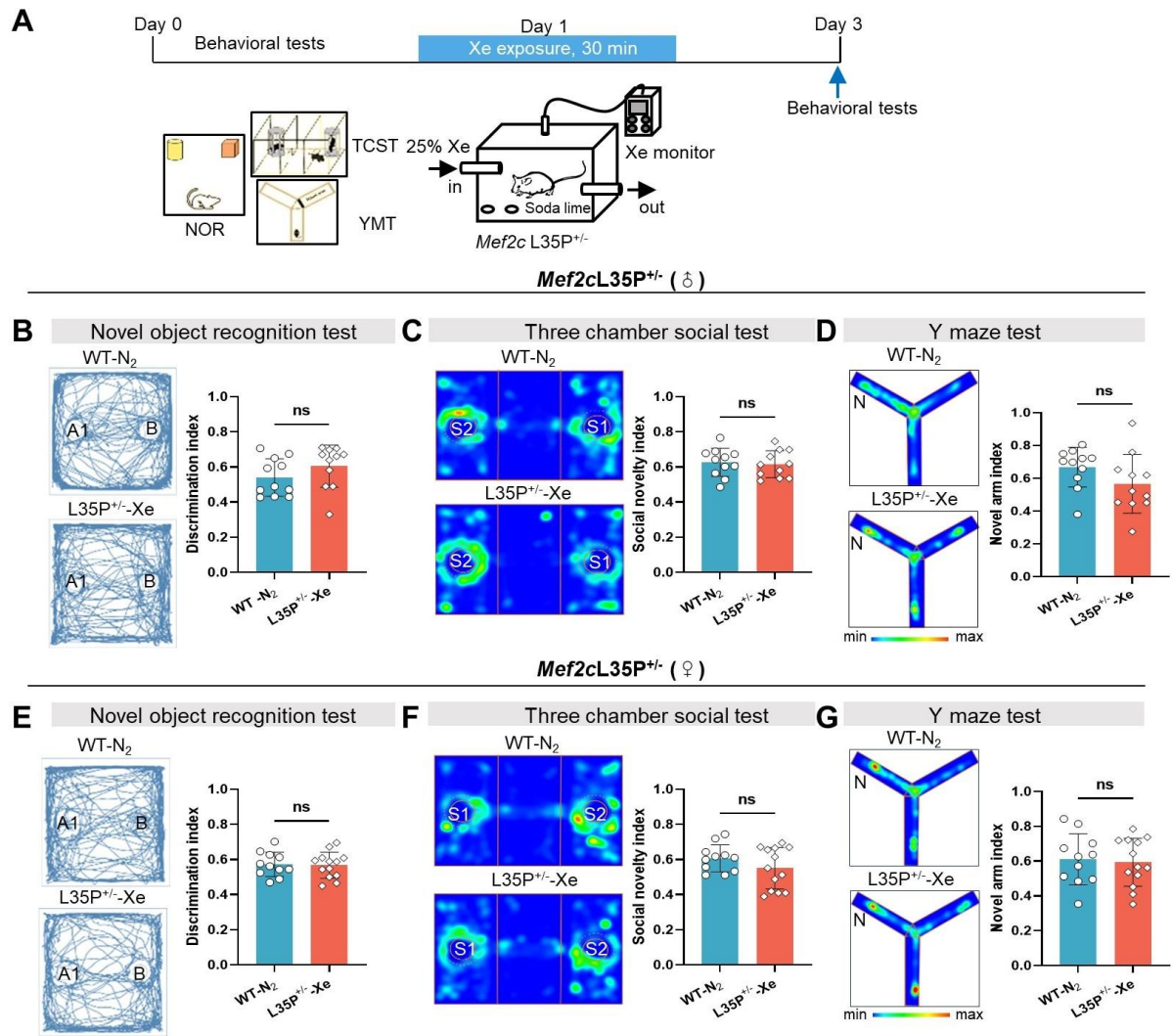

**fig. S2. Short-term Xe exposure improved autism-like behaviors in L35P<sup>+/-</sup> male and female mice.** (A) Experiment timeline of xenon (Xe) exposure and behavioral tests including novel object recognition test (NORT), three-chamber social test (TCST) and Y-maze test (YMT). (B–D) Behavioral performance in male mice 2 days after Xe exposure. Representative movement traces or heatmaps (left) and quantification of discrimination index in NORT (B), social novelty index in TCST (C), and novel arm preference index in YMT (D). (E–G) Behavioral performance in female mice 2 days after Xe exposure. Representative movement traces or heatmaps (left) and quantification of discrimination index in NORT (E), social novelty index in TCST (F), and novel arm preference index in YMT (G). Data are presented as mean ± SEM. ns, not significant.

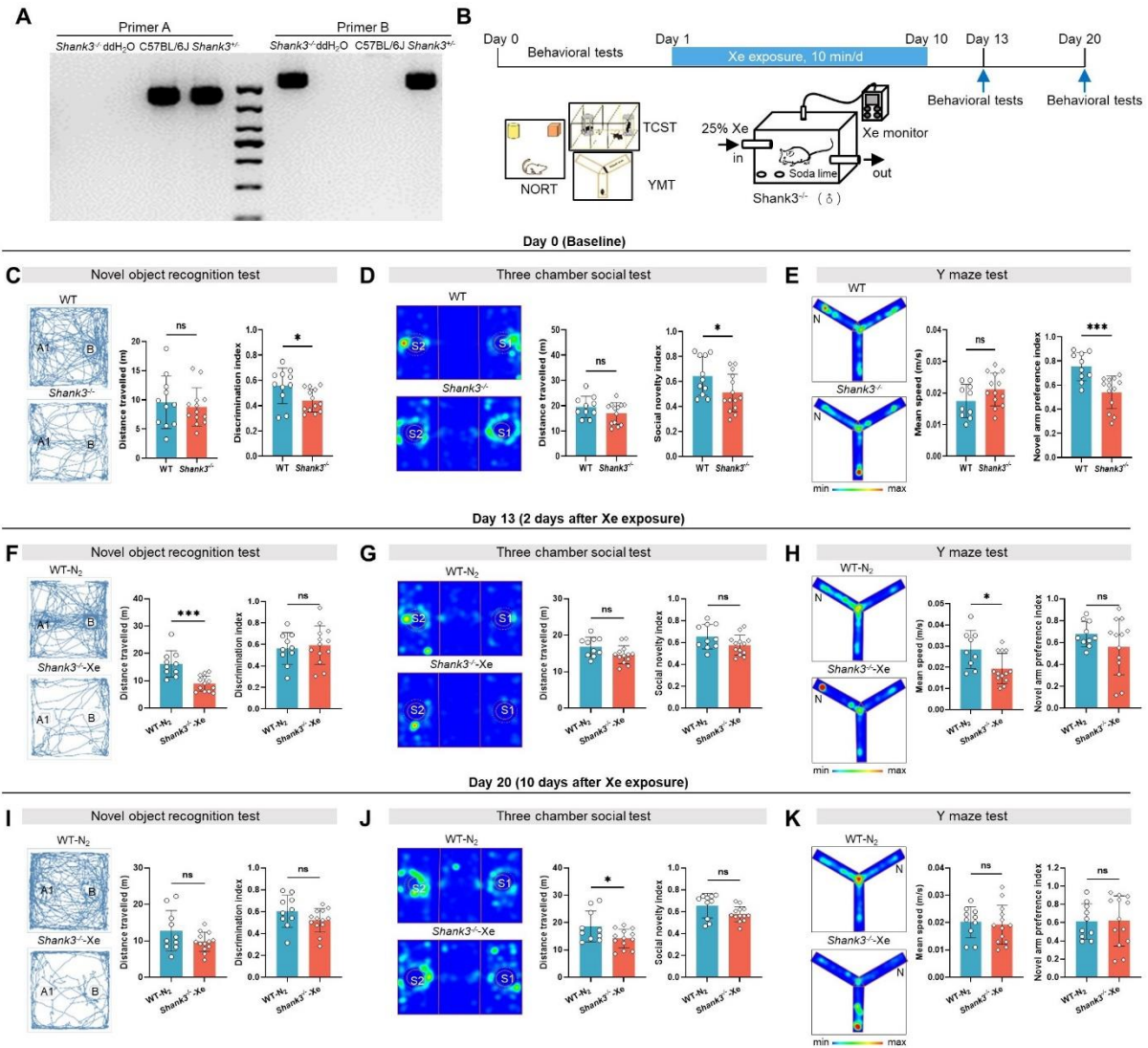

**fig. S3. Xe improved autism-like behaviors in *Shank3*<sup>-/-</sup> male mice.** (A) Genotyping (left) and Sanger sequencing (right) confirming wild-type (WT) mice and *Shank3*<sup>-/-</sup> male mice. (B) Experiment timeline of xenon (Xe) exposure and behavioral tests including novel object recognition test (NORT), three-chamber social test (TCST) and Y-maze test (YMT). (C–E) Baseline behavioral performance in *Shank3*<sup>-/-</sup> male mice without Xe exposure. Representative movement traces or heatmaps (left) and quantification of discrimination index in NORT (C), social novelty index in TCST (D), and novel arm preference index in YMT (E). (F–H) Behavioral performance in *Shank3*<sup>-/-</sup> male mice 2 days after Xe exposure. Representative movement traces or heatmaps (left) and quantification of discrimination index in NORT (F), social novelty index in TCST (G), and novel arm preference index in YMT (H). (I–K) Behavioral performance in *Shank3*<sup>-/-</sup> male mice 10 days after Xe exposure. Representative traces or heatmaps (left) and quantification of discrimination index in NORT (I), social novelty index in TCST (J), and novel

arm preference index in YMT (**K**). Data are presented as mean ± SEM. \*$P < 0.05$, \*\*$P < 0.01$,
297 and \*\*\*$P < 0.001$.

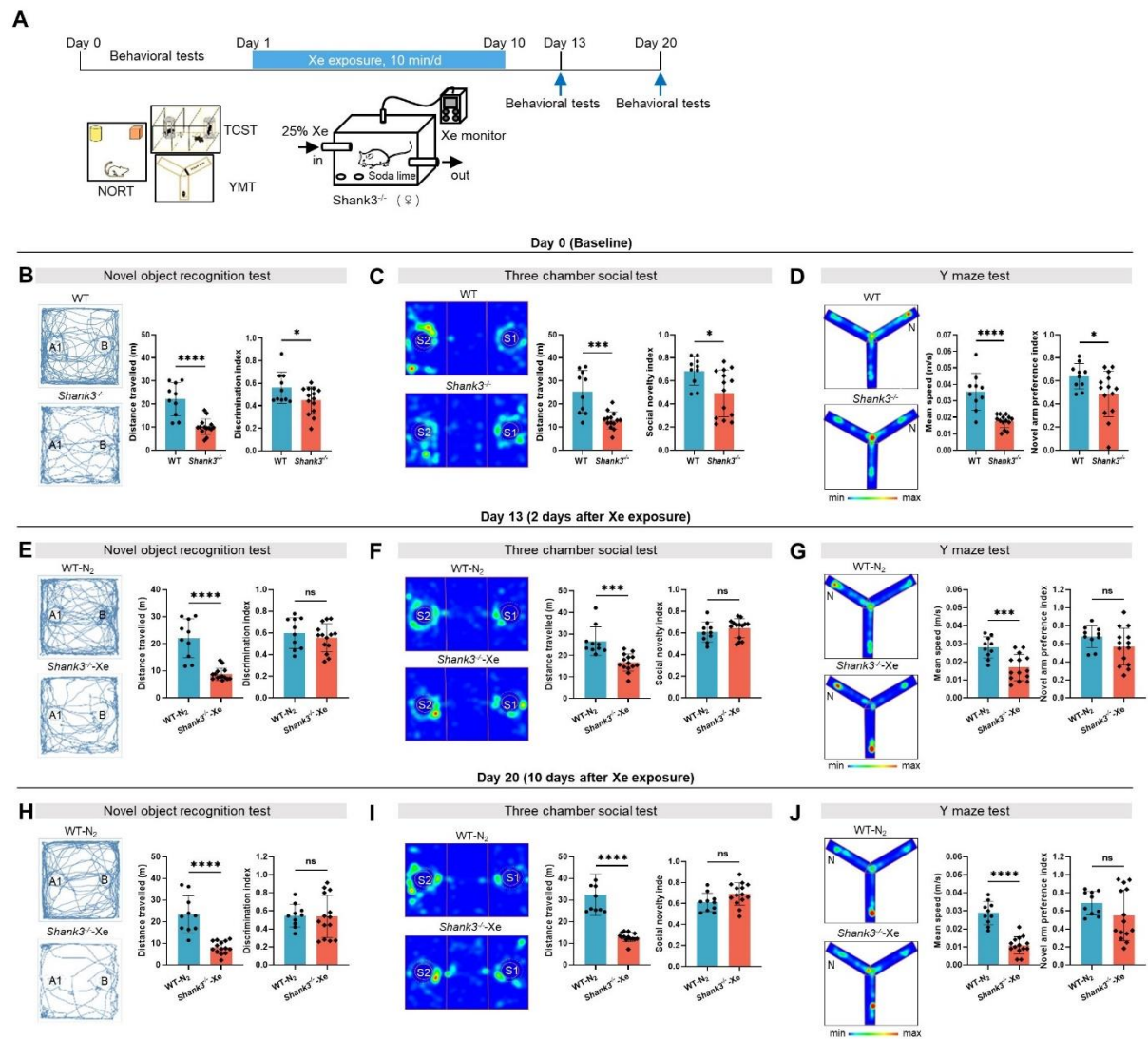

**fig. S4. Xe improved autism-like behaviors in *Shank3*<sup>-/-</sup> female mice. (A)** Genotyping (left) and Sanger sequencing (right) confirming wild-type (WT) mice and *Shank3*<sup>-/-</sup> female mice. **(B)** Experiment timeline of xenon (Xe) exposure and behavioral tests including novel object recognition test (NORT), three-chamber social test (TCST) and Y-maze test (YMT). **(C–E)** Baseline behavioral performance in *Shank3*<sup>-/-</sup> female mice without Xe exposure. Representative movement traces or heatmaps (left) and quantification of discrimination index in NORT **(C)**, social novelty index in TCST **(D)**, and novel arm preference index in YMT **(E)**. **(F–H)** Behavioral performance in *Shank3*<sup>-/-</sup> female mice 2 days after Xe exposure. Representative movement traces or heatmaps (left) and quantification of discrimination index in NORT **(F)**, social novelty index in TCST **(G)**, and novel arm preference index in YMT **(H)**. **(I–K)** Behavioral performance in *Shank3*<sup>-/-</sup> female mice 10 days after Xe exposure. Representative traces or heatmaps (left) and quantification of discrimination index in NORT **(I)**, social novelty index in

TCST **(J)**, and novel arm preference index in YMT **(K)**. Data are presented as mean ± SEM. \*$P$ <
314 0.05, \*\*$P$ < 0.01, \*\*\*$P$ < 0.001 and \*\*\*\*$P$ < 0.0001.

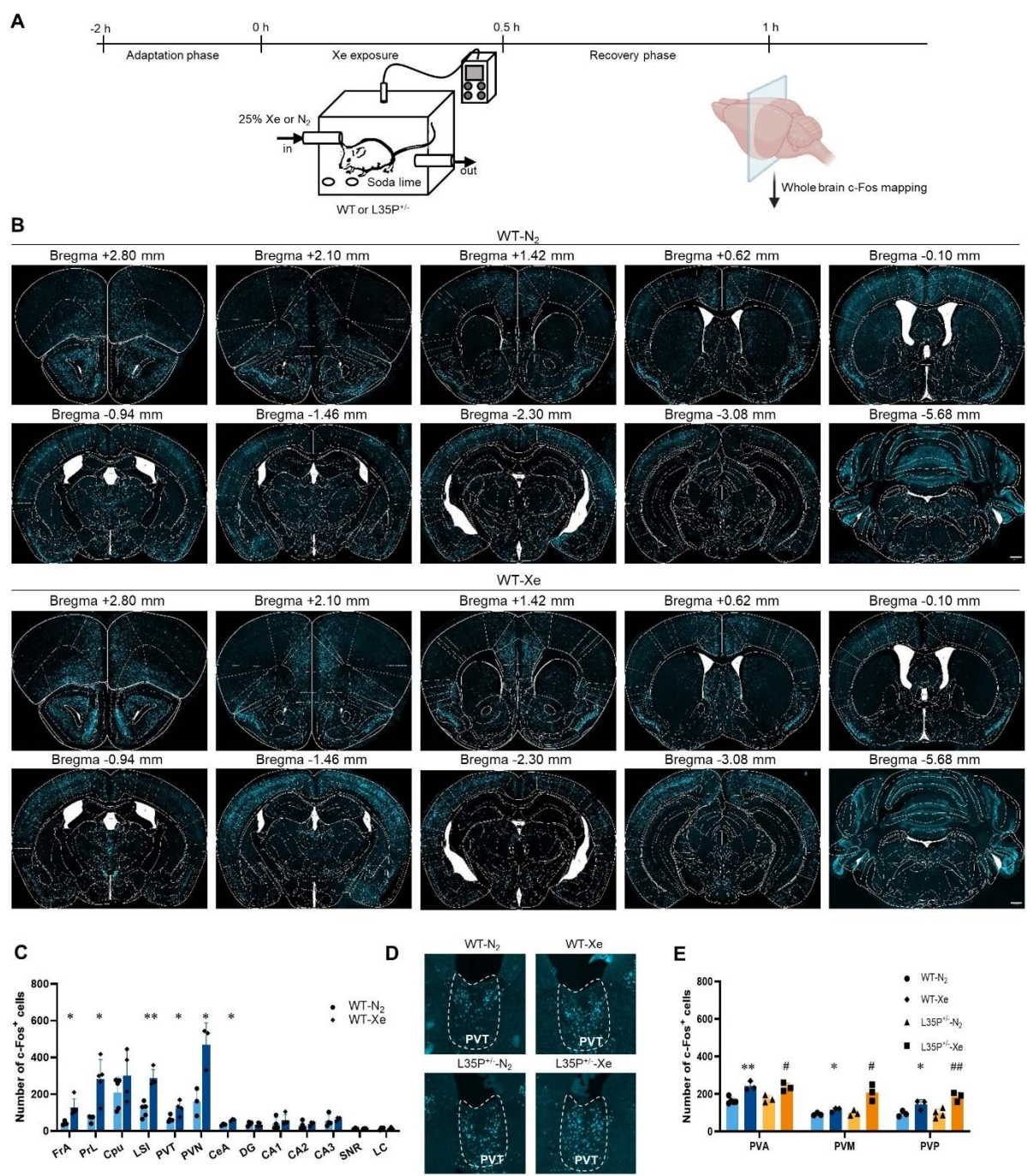

**fig. S5. Xe increased PVT neuronal activity in L35P<sup>+/</sup> mice. (A)** Experiment timeline of xenon (Xe) exposure and whole-brain c-Fos immunostaining. **(B)** Whole brain c-Fos expression in wild-type (WT) mice exposed to Xe or nitrogen (N<sub>2</sub>). **(C)** Quantification of c-Fos expression across multiple brain regions in both groups. FrA, frontal association cortex; PrL, prelimbic cortex; CPu, audate putamen (striatum); LSI, intermediate part of the lateral leptal nucleus; PVT, paraventricular thalamic nucleus; PVN, paraventricular nucleus of the hypothalamus; CeA, central amygdala; DG, dentate gyrus; CA1, field CA1 of hippocampus; CA2, field CA2 of hippocampus; CA3, field CA3 of hippocampus; SNR, substantia nigra, reticular part; LC, locus

coeruleus. (D) Representative images showing c-Fos expression in the PVT of WT and L35P<sup>+/-</sup> mice exposed to N<sub>2</sub> or Xe. (E) Quantification of c-Fos expression in the anterior (PVA), middle (PVM), and posterior (PVP) part of the PVT in WT and L35P<sup>+/-</sup> mice exposed to N<sub>2</sub> or Xe. Data are presented as mean ± SEM. \**P* < 0.05, and \*\**P* < 0.01; #*P* < 0.05, and ##*P* < 0.01.

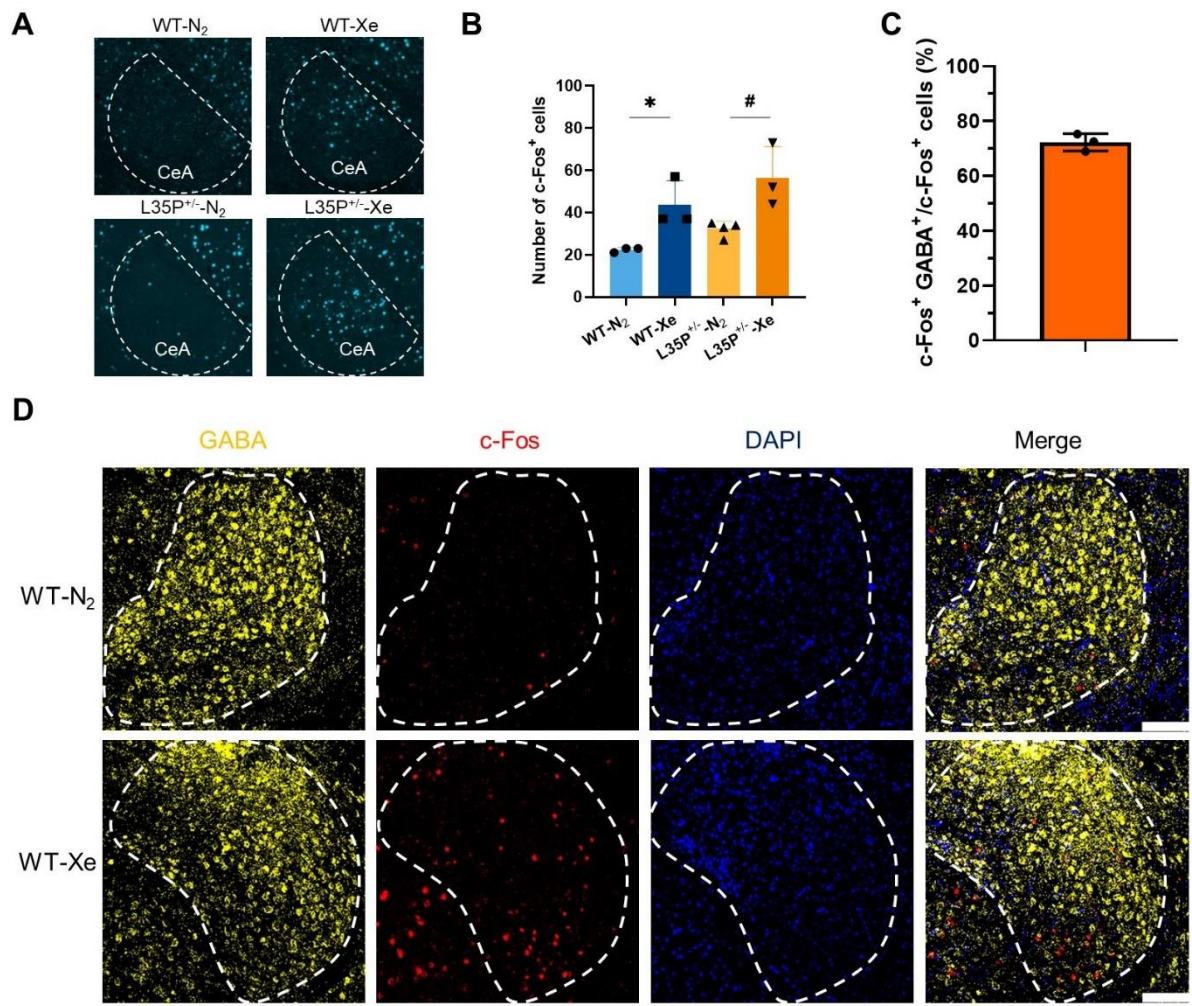

**fig. S6. Xe increased GABAergic neuronal activity in the CeA of L35P<sup>+/-</sup> mice.** (A) Representative images showing c-Fos expression in the central amygdala (CeA) of wild-type (WT) and L35P<sup>+/-</sup> mice exposed to nitrogen (N<sub>2</sub>) or xenon (Xe). (E) Quantification of c-Fos<sup>+</sup> neurons in the CeA of WT and L35P<sup>+/-</sup> mice exposed to N<sub>2</sub> or Xe. (C) Percentage of c-Fos<sup>+</sup>GABA<sup>+</sup> neurons relative to total c-Fos<sup>+</sup> neurons in the CeA. (D) Representative images showing co-localization of c-Fos and GABA in the CeA of L35P<sup>+/-</sup> mice exposed to N<sub>2</sub> or Xe. Data are presented as mean ± SEM. \**P* < 0.05, and #*P* < 0.05.
